# Antarctic photosynthesis: energy transfer and charge separation in the diatom *Chaetoceros Simplex*

**DOI:** 10.1101/2025.01.08.631881

**Authors:** Shu En Lee, Willem van de Poll, Volha Chukhutsina

## Abstract

The polar oceanic environment poses extreme challenges to photosynthetic organisms, which have evolved atypical strategies to maintain efficient photosynthesis in cold temperatures. Here, the psychrophilic diatom *Chaetoceros simplex (C. simplex)* is studied *in vivo* in the dark-adapted state using steady-state and time-resolved fluorescence methods. Our results show that all fucoxanthin chlorophyll a/c protein (FCP) antenna transfer energy to photosystem I (PSI) or photosystem II (PSII), with no detached FCPs. PSI exhibits no fluorescence of ‘red’ forms of chlorophyll (chl) beyond 700 nm in both 279 K and 77 K conditions. Despite this, it apparently has a long decay time of ∼85 ps indicating the presence of a large core-antenna supercomplex. PSII has an average lifetime of ∼500ps in open state (Q_A_ oxidized) and ∼1220 ps in closed state (Q_A_ reduced). PSII of C. simplex has kinetics that are slightly slower than temperate diatoms, suggesting slightly larger antenna. In addition, fucoxanthin (fx) molecules of FCP that absorb in the 500 - 550 nm range (fx-red) transfer more energy to PSII than fx that absorb in the blue range (fx-blue, 462 nm max absorption). A subpopulation of red-shifted, aggregated FCPs are detected at 77 K, that are active in energy transfer uphill at 279 K. Overall, our results indicate relatively larger antenna of PSI and PSII and an absence of red chls in PSI of cold-adapted species, compared to temperate species.

## 1. Introduction

The polar oceanic regions experience low temperatures, low micronutrient levels, and drastic yearly variations in photoperiod and light intensity, going from summer days of 24-hour sunlight, to winter days where the sun does not rise above the horizon (1). The Arctic and Antarctic ecosystems sequester at least 32% of global CO_2_ annually (2). In addition, the northern Arctic and southern Antarctic Oceans together constitute approximately 20% of the Earth’s surface. Polar photosynthesis is therefore of great importance to an overall understanding of photosynthesis, especially in the context of climate change (3,4). Diatoms are one of the dominant phytoplankton groups in the polar regions, and overall diatoms have been extremely diverse and evolutionarily successful at adapting to various aquatic environments on Earth (5,6). They are unicellular photosynthetic eukaryotes that can respond rapidly and flexibly to changes in light intensity of several orders of magnitude in turbulent waters.

Diatoms possess membrane-intrinsic antenna called fucoxanthin chlorophyll a/c binding proteins (FCP) that use the carotenoid fucoxanthin (fx) and chlorophyll c (chl c) as the main and accessory light harvesting pigments, respectively. FCPs can be classified based on their sequence as Lhcf, Lhcr, Lhcx, plus several other minor subfamilies (7–9). The presence of fx allows them to absorb light across a wide range of blue-green wavelengths (460-550 nm). Diatoms are unable to perform state transitions and have either no or very little spillover mechanism to rebalance excitation energy between photosystem (PS) I and II (10–12). Diatoms do, however, have sophisticated non-photochemical quenching (NPQ) systems regulated by the diadinoxanthin/diatoxanthin (ddx/dtx) cycle and the violaxanthin (Vx) cycle, in which PSII antenna play a role (13–15). NPQ has been shown to be dependent on the Lhcx subfamily of FCPs that are structurally related to green algal LhcSR (7,16). Therefore, diatom antenna have both light-harvesting and photoprotective roles. Psychrophilic green algae have unique strategies (3) for adjusting the efficiency of light harvesting and photoprotection, hence the study of psychrophilic diatoms – from the red lineage - can shed light on the possible role of antenna in their light harvesting and photoprotective qualities in polar environments (2,3,17).

In diatoms, there is both spectroscopic and structural evidence for differences in antenna associated with PSI versus PSII. Initial characterization of FCPs came from biochemical studies on isolated complexes of FCP (not bound to any photosystem). These FCPs contain about 6 - 8 chl a : 6 - 8 fx : 2 chl c, (plus 1 or less diatoxanthin / diadinoxanthin) (10,18,19). Miloslavina et al. found no difference in the excited state decay kinetics of FCPs from *C. meneghiniana* and *P. tricornutum* (average lifetime of 4.7 ns, with individual lifetimes of 130 ps, 1.8-2.6 ns and 5.5 ns respectively), implying that such FCPs are identical in both centric and pennate diatoms (20). These FCPs form trimers and oligomers in centric diatoms, and in *C*. *meneghiniana* the trimer is also the basic subunit of the oligomer (7,21,22). In pennate diatoms only trimers are present (23). Diatoms contain many FCP genes divided into the subfamilies mentioned earlier (Lhcf, Lhcr, Lhcx and others); for example, 46 genes have been identified in *Chaetoceros gracilis (C. gracilis)* and no less than 26 in *C. meneghiniana*. (8,24)

In recent years, the structures of core-antenna supercomplexes of diatoms have been determined with cryo-EM / cryo-ET. The FCPs resolved structurally were then fitted to the product of well-known FCP genes. Such supercomplexes are labelled PSI-FCPI and PSII-FCPII respectively (10,19,25–32). These FCPs, tightly bound to the photosystems, show large variation in pigment ratios of fx : chl a : chl c bound (8,31), with some FCPs containing unique additional binding sites, and some binding sites being promiscuous for either chl a or chl c (8,31). The FCP pigment ratios are highly tuned to that FCP’s specific function in the supercomplex and the presence/absence of pigments at specific locations mediates energy transfer among adjacent FCPs or from FCPs to the core (8,18,26,29,31,32). In *C. gracilis* PSI-FCPI, which contains 24 monomeric FCPs surrounding a PSI core, contains 326 chl a, 34 chl c, 102 fx, 35 ddx, and 18 β-carotenes (31) with a molecular weight of 1.1 MDa, and is the largest PSI supercomplex ever found in any photosynthetic organism. Therefore, FCPs are now known to be much more diverse in structure and function than previously observed.

The recently published isolated *P. tricornutum* FCP structure does not agree with several observations from ultrafast spectroscopy on the isolated FCPs of *C. meneghiniana* (8). For example, spectroscopy measurements have generally detected two excitation energy transfer pathways of fx to chl a, with different timescales and efficiencies of energy transfer (80% versus 60%) (18,33). However, in *P. tricornutum* FCP structure (19), all fx to chl a distances are quite similar. Furthermore, pump-dump probe results from *P. tricornutum* FCP indicate two different pathways for EET from the S_1_/ICT state of fx to the Q_y_ state of chl a, that occur with very different lifetimes than those observed in *C. meneghiniana* (34). In *C. meneghiniana*, 2D spectroscopy shows that there should be two spatially separated chl c with quite different connections to chl a with correspondingly different timescales of energy transfer (390 fs vs 4.5 ps). Assuming Forster level of description, calculations of the expected dipole-dipole interactions in *P. tricornutum* structure indicate that both chl c should have similar rates of EET to chl a. However, the calculations assumed that all chl a have the same energy, and all chl c have the same energy, a rather severe approximation. Lastly, in *P. tricornutum* FCP dimer structure two fx (fx306 and fx307) are in close contact with chl c, but far from any chl a (19). This would imply that some energy transfer from fx to chl c occurs, yet this has never been detected spectroscopically (8,10). Gelzinis et al (8) attempted to build a theoretical *C. meneghiniana* FCP model consistent with spectroscopic evidence and arrived at a trimeric structure. Later on, a trimer loosely associated with PSII was isolated and its structure was found to be in good agreement with the model of Gelzinis and co-workers (32). Therefore, EET pathways are highly species specific.

The two different EET pathways from fx to chl a in *C. meneghiniana* have been ascribed to two spectrally distinct fx. Fx can be strictly described as fx-blue (λmax = 463 nm), fx-green (λmax = 492 nm), or fx-red (λmax = 500 – 550 nm), although the broad absorption peaks of fx means that their absorptions necessarily overlap (35–38). Fx-red and fx-blue have different efficiencies of energy transfer to chl a (80 % vs 60 % if fx-red and fx-blue are defined as absorbing > 510 nm and < 510 nm respectively) (18,33,39). There are contradictory results regarding the roles of these two fx in energy transfer to PSI or PSII (13,39). Chukhutsina et al. 2013 and Chukhutsina et al. 2014 (13,40) demonstrated that in *C. meneghiniana* most antenna are connected to PSII, with markedly lower PSI fluorescence when fx is excited. Furthermore, they concluded that fx-red mostly transfers energy to PSII, while fx-blue transfers proportionally more energy to PSI. In Zhao et al (32) an isolated FCP trimer in *C. meneghiniana* was enriched in fx- red and loosely associated with PSII. In contrast, other studies paint a different picture. In *C. meneghiniana*, the FCPb absorption spectrum appears to show more fx-red and less fx-blue content compared to FCPa (41). Additionally, solation and biochemical characterisation of FCPb indicated that its sole component is Fcp5 which was loosely associated with PSI (42). Therefore, this is also an area of considerable interest.

Despite the abundance of literature on isolated complexes, there are relatively few studies on intact cells. Studying intact cells can provide more comprehensive physiological picture, since isolation and preparation conditions can influence the size and composition of complexes. Miloslavina et al (20) arrived at a kinetic model for closed-state diatoms with an average PSI lifetime of ∼51 ps and average PSII lifetime of ∼395 ps for *C. meneghiniana* and ∼100 ps and ∼431 ps for *P. tricornutum*. In their target model, PSII is described by one excited state compartment of antenna and reaction centre (Ant/RC) and two radical pairs (RPs). PSI is described by one Ant/RC, one RP and one “red” compartment assigned to red-shifted chls. Higher plants characteristically require two red compartments to fully describe PSI kinetics. However, in diatoms the evidence for red chls is equivocal, as some studies have identified red chls while others have not (43,44). Yokono et al (45) studied the fluorescence decay of cells of four different diatom species at 77 K and concluded that all had red chl species in PSI. Chukhutsina et al (13) observed the presence of red chls in *C. meneghiniana* PSI emitting at around 717 nm, agreeing with Veith et al (42). Such red chls, if present, may be located in the antenna rather than the core: Ikeda et al (43) did not detect red chl species in isolated PSI core of *C. gracilis*. Moreover, a growing body of evidence indicates that a specific red-shifted FCP contains red forms, appearing under conditions of chromatic acclimation to red light (30,46–50). In the Antarctic green alga *P. crispa*, red chls transferring energy to PSII were suggested to be an adaptation to far-red light conditions (51). However, in the single-celled *Chlamydomonas priscuii* (*C. priscuii*), another polar green alga, only minimal red fluorescence beyond 700 nm was seen (52). The presence or absence of red forms in polar organisms is hence also an area of significant interest.

This paper aims to explore the role of antenna and photosystems in modulating energy transfer and charge separation in the Antarctic centric diatom *Chaetoceros simplex* (*C. simplex*) in the dark-adapted state, mainly through global and target analysis of time-resolved spectra. Experiments at 279 K and 77 K utilise different excitation wavelengths to preferentially excite cores or antenna. Overall, we determine how antenna are connected to the photosystems and how *C. simplex* compares to other species in its kinetics of energy transfer and charge separation.

## 2. Materials and Methods

### 2.1 Cell cultures and absorption spectrum

The diatom *C. simplex* isolated from the coastal waters of Prydz Bay, Antarctica (Australian National Algal Culture Collection, Bacillariophyceae, CS 624, ANACC, 3 - 5 µm) was grown in 250 ml cell flasks (Cellstar) at 4°C in low white light of 10 - 15 µE (spectrum in Supplementary Fig. 1). Cells were suspended in F/2-enriched sea water (53,54), and exposed to a 12 h light: dark cycle. Cells were acclimated to these conditions for months and were only harvested for experiments during the exponential phase. The absorption spectrum of cells was measured regularly with an Agilent Cary 4000 spectrometer every 2 - 3 weeks to observe any changes in pigment expression in the exponential phase (no changes were observed).

### 2.3 Steady-state fluorescence measurements

Steady state emission spectra were obtained at 77 K and 279 K (6°C) with a Jobin Yvon Fluorolog. Different excitation wavelengths were used to excite different pigments, at 400 nm, 440 nm, 467 nm, 520 nm and 550 nm. A slit width of 5 nm and 0.5 nm was used for excitation and emission respectively, with an integration time of 1 s. An O.D. titration was performed by measuring the emission of *C. simplex* samples at different O.D.s (Supplementary Fig. 2)

### 2.4 Time-correlated single photon counting (TCSPC)

Time-correlated single photon counting (TCSPC) measurements were performed at 279 K with a FluoTime 200 (PicoQuant) using a 440 nm or 467 nm laser diode at a frequency of 2.5 MHz. The excitation spot was ∼200 µm in diameter. Photons emitted from the sample were detected with a Hamamatsu MPC-PMT detector. The decay of pinacyanol iodide in methanol, which has a lifetime of 6 ps, was measured at 680 nm detection both before and after the experiment (FWHM of 88 – 92 ps) (Supplementary Fig. 4) (55). Using the pinacyanol decay trace the IRF signal was deconvoluted from the sample decay. A quartz cuvette with a 1 cm path length was used and the sample was constantly refreshed with a magnetic stirrer. For each measurement, the sample was concentrated to an absorbance of approximately 0.25 at the Q_y_ peak of chl a.

The samples were measured in two states, “open” and “closed”, corresponding to oxidised and reduced Q_A_ in photosystem II respectively. Open state was achieved by dark-adapting the sample for 15 minutes. Closed state was achieved by the addition of 100 µM DCMU and 1mM hydroxylamine, followed by illumination for 2 - 3 minutes under weak (24 uE) light (Supplementary Fig. 3). Closure of PSII RCs was deemed successful when the level of Fo rises to the dark-adapted Fm level measured on a Dual-PAM (Dual-PAM 100, Walz, Germany). For the ‘open’ state, power titrations were performed varying the laser power to ensure that samples were truly ’open’ (Supplementary Fig. 6).

Decay traces were then collected using a wave step of 5nm from 660 nm to 730 nm. A time step of 4 ps was used and traces were measured up to an 8 ns time window. At the end of the experiment, a check was performed to ensure that the sample was stable for the duration of the experiment (Supplementary Fig. 5). A comparison of decay traces from different samples indicated some variability in lifetimes (Supplementary Fig. 7). Decay traces were then fitted to multi-exponential decay functions using the program TRFA Advanced (SSTC) (56). The lifetimes τ_k_ were linked globally across the different detection wavelengths, but amplitudes were allowed to vary. Quality of the fit was judged by the reduced chi-squared parameter (χ2), the plot of weighted residuals and their autocorrelation function (Supplementary Fig. 8).

TRFA fits fluorescence decay traces using the following Equation 1:

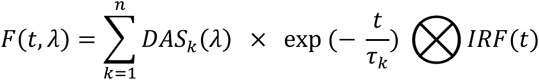

where *DAS_k_* is the decay-associated spectrum and *τ_k_* is the related decay lifetime.

Target analysis was done using the program Target 3.1. Unlike global analysis, which assumes independent, unconnected compartments undergoing parallel decay, target applies a specific kinetic model where transitions between compartments are described by rate constants. The concentration profile of components over time can be obtained analytically (57).

### 2.5 Streak camera measurements

At 279 K, the sample was excited at 400 nm, 475 nm and 520 nm. At 77 K, it was excited at 400 nm, 475 nm, 520 nm and 550 nm. A band pass filter (+- 5 nm) was used to cut out light at all other wavelengths. Sample was measured in the “closed” state and was concentrated to an optical density of <0.5 at the excitation wavelength. At 279 K a quartz cuvette of path length 1 cm was used and the beam was focused onto the cuvette edge directly facing the detector. At 77 K, a glass capillary of path length 1 mm was used. The laser power was around 100 µW at 279 K and 100 – 200 µW at 77 K. Fluorescence emission from the sample was focused into a spectrograph (Chromex 250IS, 50 grooves mm−1) and the output light from the spectrograph was then focused onto the streak camera (Hamamatsu C5680, Shizuoka, Japan).

For each dataset, 30 streak images were recorded, with each image being an analog integration of 60 exposures with exposure time of 1 s each. Before analysis the images were averaged into a single dataset and then corrected for background signal and wavelength sensitivity of the detector. Two different time windows were used for measurement: 0.45 ns (time range 2) and 1.5 ns (time range 4).

Streak images were analyzed using Glotaran software for global analysis (58). The IRF was modelled in silico, using a single Gaussian whose width could vary during fitting. The FWHM of the IRF was determined to be ∼8 ps for time range 2 and ∼23 ps for time range 4. Quality of fit was judged by the singular value decomposition of the residuals matrix in time (Supplementary Fig. 9 and Fig. 10). Glotaran fits fluorescence decay traces using Equation 1 mentioned above in Section 2.4. The steady-state emission spectrum and the total fluorescence spectrum at time zero were reconstructed (Supplementary Fig. 12 and 13), demonstrating agreement with emission spectrum results in section 3.1.

## 3. Results

### 3.1 Emission spectra

We measured the steady-state emission spectra of cells at 279 K (6°C, corresponding to the temperature in *C. simplex*’s native habitat) and 77 K (Figure 1). At 279 K, there is a broad main peak centering around 680 nm that is mostly associated with PSII. There is no clear emission from PSI in the red region (>700 nm). At 77 K this main PSII-associated peak red shifts to 686 nm and a shoulder’ appears at 695 nm and slowly tails off ending at around 715 nm. Again, there is no clear red PSI emission at >700 nm, unlike that seen in *C. meneghiniana* at ∼720 nm (40). At both 77 K and 279 K there is a vibronic band from 720 - 760 nm due to chlorophylls. Furthermore, regardless of the excitation wavelength, ranging from 400 nm to 550 nm, emission spectra are always identical, indicating that there is no detached antenna. All FCPs are associated with either one or both PS, and it is possible that all FCPs in *C. simplex* are tightly associated with cores.

**Figure 1.**
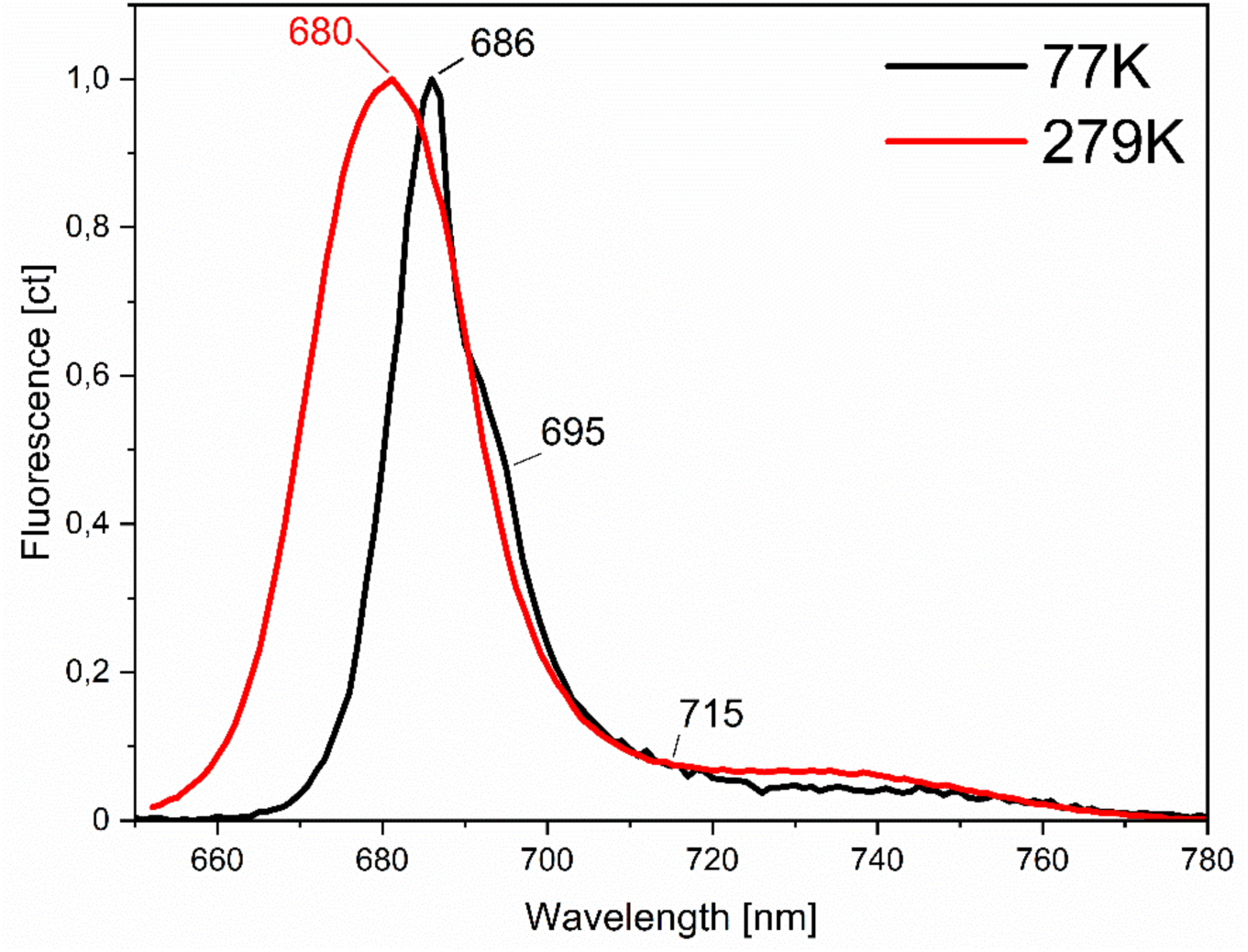
Steady-state emission spectra of C. simplex at 279 K and 77 K, normalised to maximum emission. Excitation wavelengths used: 400 nm, 440 nm, 475 nm, 520 nm, 550 nm. Since emission spectrum did not vary with excitation wavelength, for simplicity of presentation, only one emission spectrum for each temperature is presented.

At 279 K, our emission spectrum not only resembles that of other diatoms at room temperature (RT), but also green algae and plant thylakoids, where there is only one main peak at 680 nm arising from PSII (48,59,60). Our 77 K spectrum somewhat resembles that of *C. meneghiniana* in Kansy et al (61), which has a maximum at 685 nm (assigned to PSII antenna) and a shoulder that begins around 700 nm (assigned to PSI). In our case, however, the shoulder begins earlier, at 696 nm. In other studies diatom PSI emits as a very distinct peak around 720 nm (40,62). Our results therefore hint at a less red PSI and that most of the antenna are bound to the photosystems.

### 3.2 TCSPC results at 279 K

We performed time-correlated single photon counting (TCSPC) experiments where excitation wavelengths of 440 nm and 467 nm were used and time resolved fluorescence emission was recorded in the range 660 - 730nm. While 440 nm excitation is selective for chl a, 467 nm favours chl c and fx(-blue) absorption, and therefore excites more antenna relative to cores (but note that it is not possible to excite cores completely selectively due to the presence of chl a in FCPs).

Global fitting of the time-resolved fluorescence decay traces revealed four exponential components in all cases. The traces were normalized to the area of the total fluorescence spectrum at time zero for each dataset. The EET character (positive to negative amplitude) of the fastest component is consistent across all replicates and is corroborated by our streak data which can give an accurate estimation of its lifetime. However, since this lifetime lies beyond the time resolution of our TCSPC setup, only positive components are shown in Figure 2. In both open and closed state, the decay associated spectra (DAS) shape and amplitudes obtained are the same within error (+- 12%). However, the average PSII lifetime is increased upon 467 nm excitation.

**Figure 2.**
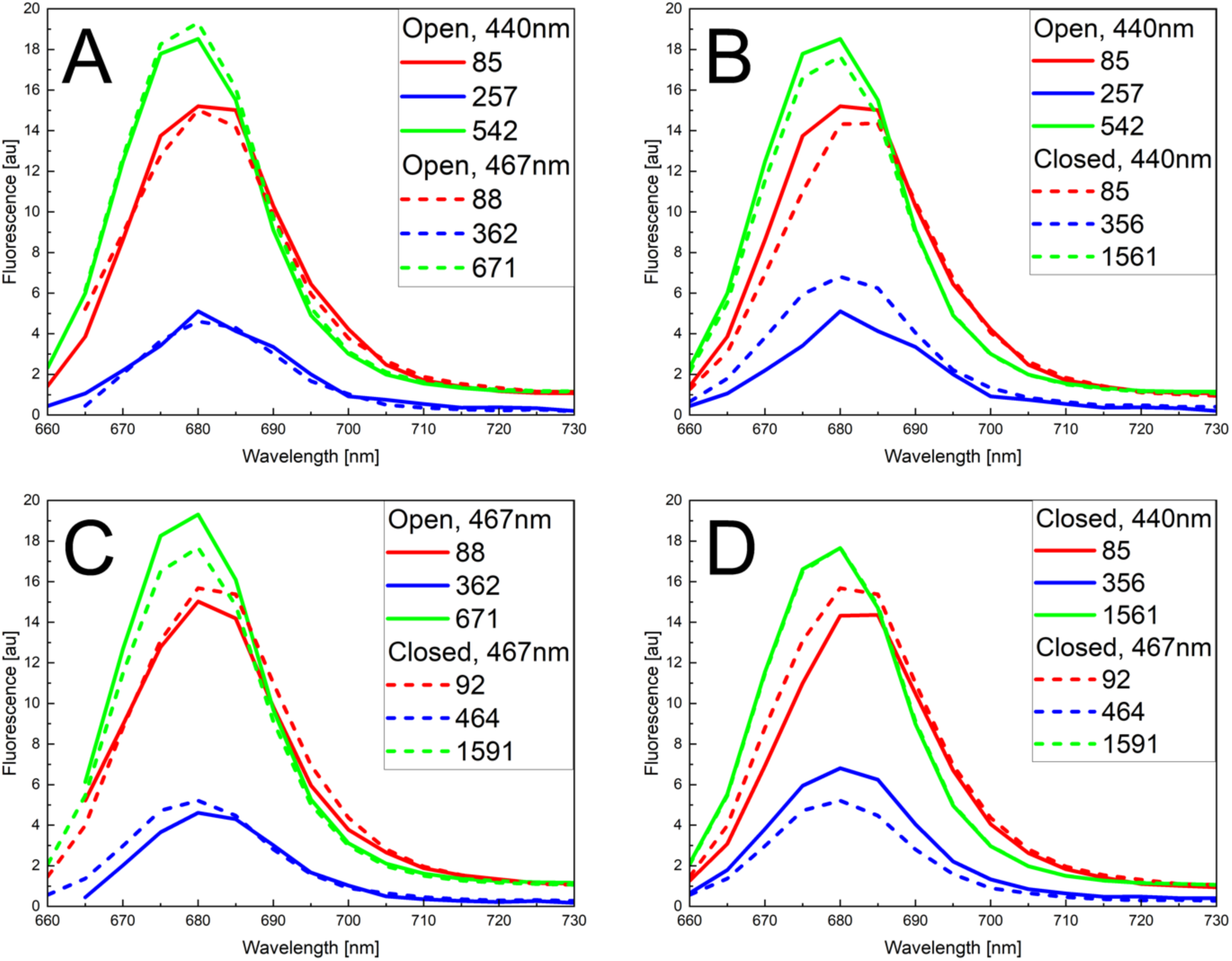
Decay Associated Spectra (DAS) of C. simplex from TCSPC experiments at 279 K. For comparison, the area of the total fluorescence spectra at t = 0 (sum of the corresponding DAS) are normalized to each other. Lifetimes are given in ps. (A) Comparison of open-state fits at 440 nm and 467 nm excitation. (B) Comparison of closed-state fits at 440 nm and 467 nm excitation. (C) Comparison of open-state and closed-state fits at 440 nm excitation. (D) Comparison of open-state and closed-state fits at 467 nm excitation. Quality of fit data is included in Supplementary section VI.

In our data, PSI is associated with a ∼85 ps component peaking at 685 - 690 nm with a spectrum that is red shifted compared to other components. This corresponds to the average excitation trapping time in PSI. In the open state, PSII is associated with two longer components of ∼240 ps and ∼600 ps peaking at ∼ 680 nm. The average PSII lifetime is 480 – 520 ps, which is much longer than the lifetime of PSII in plant thylakoids when 4 trimers are present per PSII RC (330 ps). The average lifetime is equal to the sum of the migration time τ_mig_ and the trapping time τ_trap_, τ_avg_ = τ_mig_ + τ_trap_. The increase in average PSII lifetime in open state upon 467 nm excitation can therefore indicate the increase in average migration time (Δ τ_mig_) for excitation energy transfer (EET) from antenna to PSII RC when fx-blue/chl c is excited (63). This difference is ∼ 30 – 50 ps across different datasets, which is much longer than in *C. meneghiniana* (∼13 ps in (40)) and wild type *Arabidopsis thaliana* (plant) thylakoid membrane with 4 LHCII trimers per PSII core dimer when chl b is excited instead of chl a, also ∼13 ps (63).

In closed state, photochemical quenching is blocked, and the kinetics of PSII change while those of PSI remain unaffected. The rate of PSII charge separation decreases while that of charge recombination increases, so PSII fluorescence increases (64). Therefore, the average lifetime and the average PSII lifetime become much longer. The overall average lifetime at 440 nm excitation increases from ∼350 ps in open state to 780 ps while average PSII lifetime goes from ∼500 ps in open state to ∼1200 ps in closed state (Table 1). Both PSII lifetimes increase upon moving from open to closed state, although the difference is smaller for the shorter PSII component (∼100 ps difference) than the longer PSII component (∼1ns difference). This contrasts with the situation in plants where both lifetimes are significantly longer in closed state (65,66).

**Table 1.** Average lifetimes and average PSII lifetimes calculated from 279 K data.

|  | Average lifetimes | PSII : PSI area |
| --- | --- | --- |
| $\tau, ps$ | Open state – 440 nm<br>(TCSPC) | $1.4 \pm 12 \%$ |
| $\tau_{avg}$ | 316 | |
| $\tau_{psii}$ | 482 | |
| | Open state – 467 nm<br>(TCSPC) | $1.4 \pm 12 \%$ |
| $\tau_{avg}$ | 333 | |
| $\tau_{psii}$ | 519 | |
| | Closed state – 440 nm<br>(TCSPC) | $1.6 \pm 12 \%$ |
| $\tau_{avg}$ | 784 | |
| $\tau_{psii}$ | 1223 | |
| | Closed state – 467 nm<br>(TCSPC) | $1.4 \pm 12 \%$ |
| $\tau_{avg}$ | 801 | |
| $\tau_{psii}$ | 1338 | |
|  | Closed state – 400 nm<br>(streak) | 1.6 |
| $\tau_{avg}$ | 688 | |
| $\tau_{psii}$ | 1053 | |
|  | Closed state – 475 nm<br>(streak) | 1.85 |
| $\tau_{avg}$ | 706 | |
| $\tau_{psii}$ | 1043 | |
|  | Closed state – 520 nm<br>(Streak) | 2.0 |
| $\tau_{avg}$ | 821 | |
| $\tau_{psii}$ | 1188 | |

### 3.3 Streak measurements at 279 K

Streak camera measurements were performed at 279 K in the closed state and 77 K. At both temperatures, excitation wavelengths of 400 nm, 475 nm, and 520 nm were used, while 550 nm excitation was additionally used at 77 K. As a rough approximation based on *C. meneghiniana* FCP absorption spectrum decomposed into the contribution of individual pigments (35), at 475 nm both fx and chl c are excited. At 520 nm, only fx experiences significant excitation and fx-red is excited more than fx-blue. At 550 nm, only fx-red is excited.

In the 279 K data 4 components are sufficient to fit the decay trace (Figure 3). There is a fast EET component (17 – 30 ps), an 80 - 90 ps component associated with PSI that peaks at ∼685 nm, and two longer components of 450 – 550 ps and 1.6 ns respectively, associated with PSII and peaking at ∼678 nm and ∼682 nm respectively. The ns component is not well resolved by the time window of the streak setup and was thus fixed to 1.6ns, the lifetime of the long PSII component obtained in our TCPSC results in closed state.

**Figure 3.**
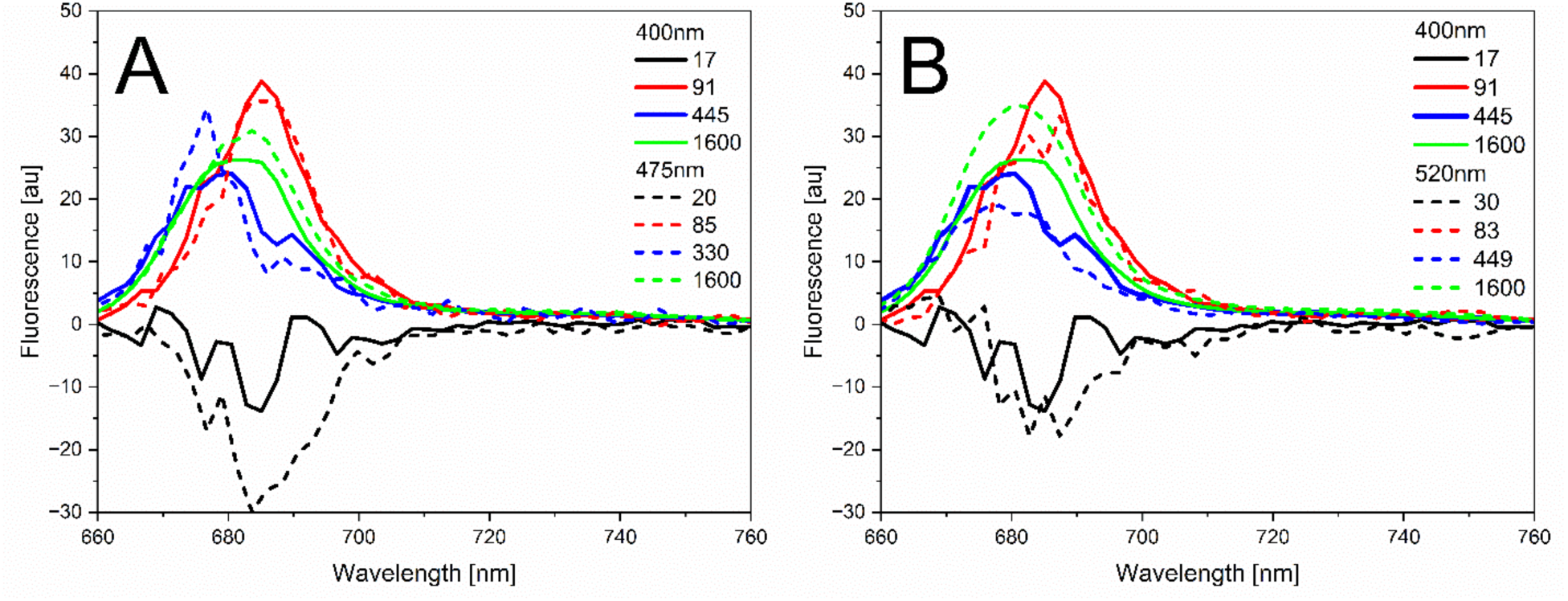
Global fitting of streak data at 279 K. (A) Comparison of 400 nm (solid lines) and 475 nm (dashed lines). (B) Comparison of 400 nm (solid lines) and 520 nm (dashed lines). For comparison, the area of the total fluorescence spectra at t = 0 (which equals the sum of all DAS) are normalized to each other. Quality of fit data is included in Supplementary section VI.

Table 1 compares the average lifetimes and average PSII lifetimes of all 279 K data, as well as the ratio of PSII to PSI areas. Since the area under the DAS scales linearly with the effective absorption cross section of that photosystem, our results at 400 nm and 440 nm show a ∼1.6 : 1 ratio of PSII : PSI areas reflecting that there are more pigments associated with PSII than PSI. This ratio increases in the order 400 nm < 475 nm < 520 nm: this suggests that PSII antenna possess longer wavelength absorbing forms of fx. In our TCSPC results, this increase in PSII : PSI area is not seen in our TCSPC results upon fx/chl c excitation; this could be due to the lower spectral resolution of TCSPC data (5 nm) compared to streak (2.7 nm).

### 3.4 Streak measurements at 77 K

### At 77 K, *C. simplex* was excited at 400 nm, 475 nm, 520 nm and 550 nm using a streak camera setup. Global fit of the streak data resolved 4 DAS (Figure 4), and the quality of fit is indicated in the residuals (Supplementary Fig 9). At low temperatures, PSII and PSI emission shifts to longer wavelengths and the emission from low energy pigments increases (67). In plants and green algae, PSI exhibits a nanosecond component at 77 K (68). However, in diatoms the contribution of the ns component is negligible or absent (13,43). In our data, the lifetimes and DAS shapes are similar for all excitation wavelengths; however, their amplitudes and peak wavelengths differ. While a 4- component fit of the 550 nm dataset is presented here in comparison with other excitation wavelengths, fitting an additional (fifth) component improved the residuals substantially and this is presented in Supplementary Material Section XI

In the 4 component DAS, the fastest component (16 – 19 ps) has a positive peak at 675 nm and a negative peak around 686 nm. The positive to negative character indicates it represents an EET process. The amplitude of the negative peak is more pronounced upon antenna excitation at 475 nm and 550 nm. In addition, the negative peak of the EET component occurs at the same wavelength (687 nm) as the positive peak of the third DAS (340 – 440 ps). The latter is assigned mostly to PSII trapping (see below); hence we assign the fast component to EET between FCPs and/or from FCPs to PSII.

**Figure 4.**
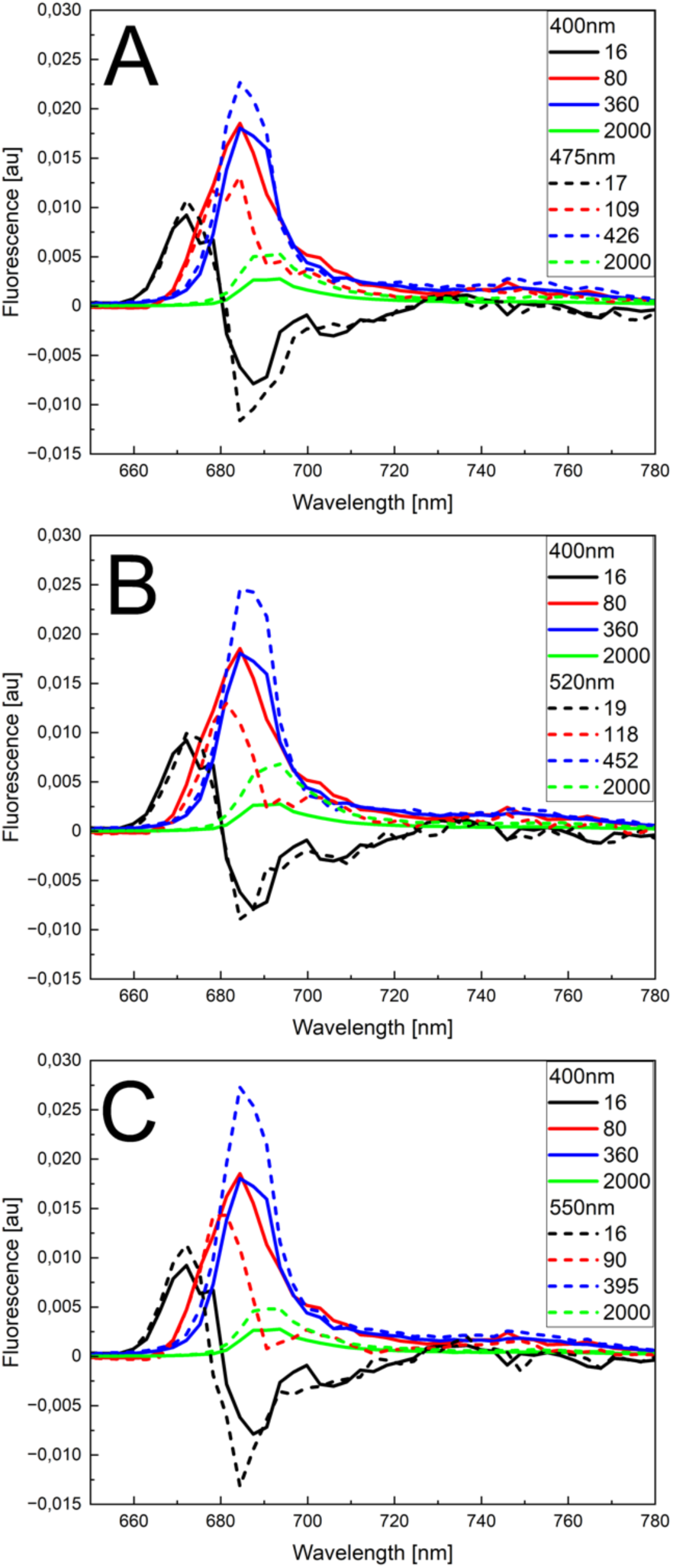
DAS of C. simplex at 77 K. (A) Comparison of 400 nm and 475 nm excitation. (B) Comparison of 400 nm and 520 nm excitation. (C) Comparison of 400 nm and 550 nm excitation. For comparison, the area of the total fluorescence spectra at t = 0 (which equals the sum of all DAS) are normalized to each other. The total fluorescence spectrum and reconstructed steady-state spectrum are published in the Supplementary Material Section VII. Quality of fit data is included in Supplementary section VI.

The second DAS (80 – 120 ps) can be decomposed into two separate Gaussian components (Figure 5). This reveals two peaks; one higher peak at ∼680 nm plus a lower, broader peak at ∼700 nm. While the former could originate from slowly equilibrating FCPs, the latter likely originates from PSI and has a rather small area. Furthermore, amplitude of the PSI peak is in the order 550 nm < 520 nm < 475 nm < 400 nm. Comparing the absorption spectrum of PSI core at 77 K to that of FCP (37,69), FCP appears to absorb relatively more energy at 475 nm than the PSI core. Assuming the stoichiometry of PSI-FCPI in *C. gracilis* (24 FCP monomers per PSI core), approximately 88 % of energy absorbed at 475 nm is due to FCP, with the remaining 12 % due to absorption by PSI core. This therefore shows that the increased PSI amplitude at 475 nm is not simply due to increased PSI core excitation. Hence, fx-blue transfers relatively more energy to PSI than fx-red. Overall, the 2^nd^ DAS resembles one in Chukhutsina et al (13) where a ∼140 ps component was associated with FCP and PSI, peaking at 680 nm and 705 nm respectively.

**Figure 5.**
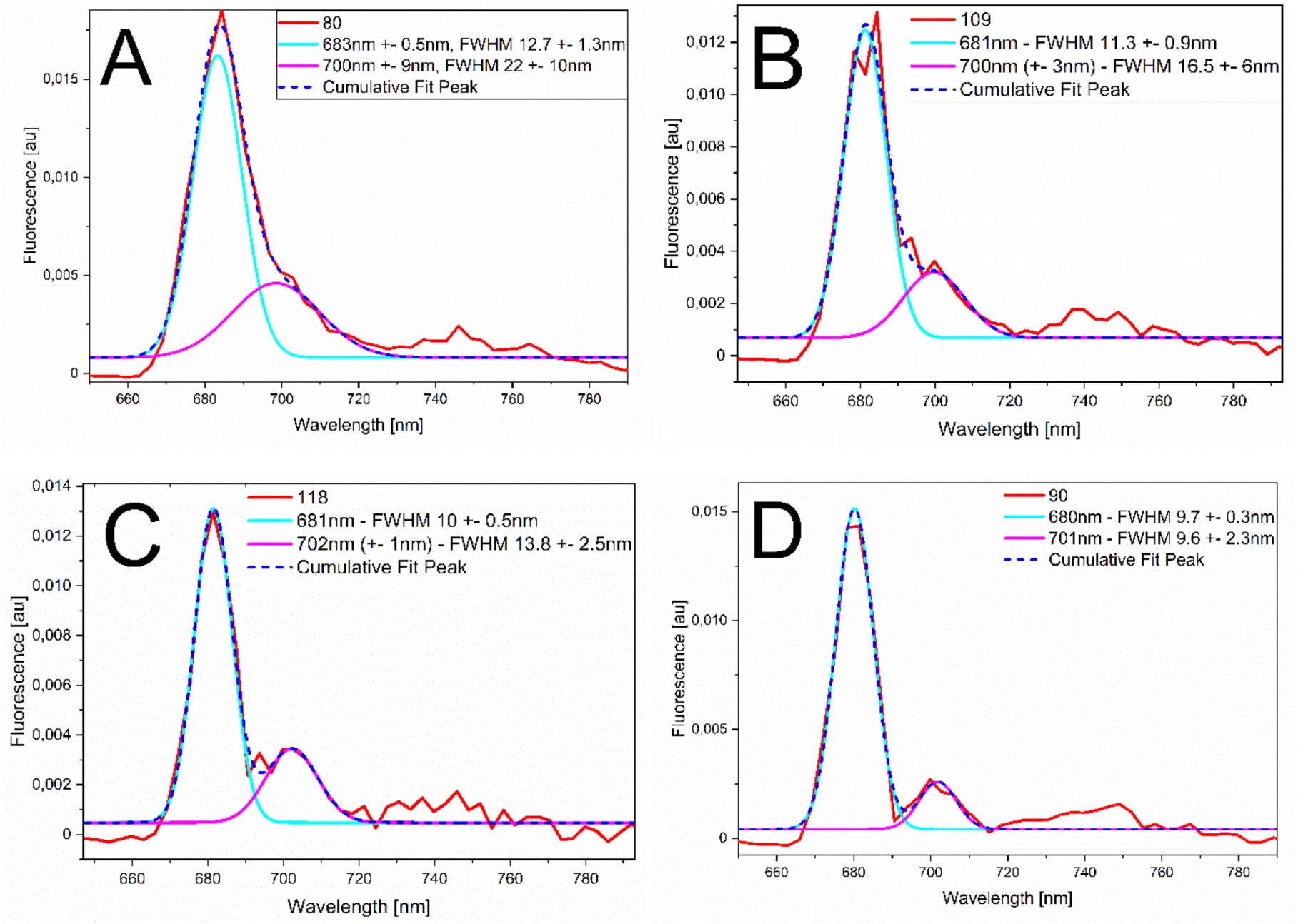
Gaussian decomposition of the 2^nd^ DAS at 77 K. Red indicates the shape of the DAS of the 2^nd^ component. (A) 400 nm, (B) 475 nm, (C) 520 nm, (D) 550 nm. The first gaussian fit is indicated in cyan and the second gaussian in magenta. FWHM – full width half maximum. (+-) indicates the uncertainty. The sum of the two gaussian fits is plotted in dark blue (dashed lines).

The third DAS (340 – 440 ps) peaking at ∼687 nm mostly originates from PSII trapping. The amplitude and relative area of this component is increased upon fx excitation, in the order 550 nm > 520 nm > 475 nm. This indicates that fx-red transfers significant energy to PSII, proportionally more than fx-blue, agreeing with our 279 K streak results.

The fourth DAS (green) with a broad, flat peak centered at 692 nm has increased amplitude upon antenna excitation, in the order 520 nm > 475 nm > 550 nm > 400 nm. This ns component lies outside the time window resolved by streak; however, multiple independent fits converged at 2 ns, hence we fixed this component at 2 ns during fitting. In *C. simplex*, the long ns component represents some red shifted, aggregated FCPs that are also seen also in *C. meneghiniana* (13,70). There is thus heterogeneity in the FCP population at 77 K. Non-aggregated FCPs emit around 670 – 675 nm, while aggregated FCPs have a lower fluorescence yield and a red shift of the fluorescence maximum to ∼690 nm (13). The aggregated red FCPs likely do not transfer energy to cores at 77 K but do transfer energy at 279 K. While the red FCPs seen upon 550 nm excitation likely mainly transfer energy to PSII at 279 K, the additional red FCPs seen at 475 nm and 520 nm excitation may also transfer energy to PSI, since their amplitude increases while the amplitude of the third/PSII component decreases. Studies on *C. meneghiniana* have found that the size of this red FCP subpopulation is different for FCPa and FCPb (trimer and oligomer respectively) and that the functional significance of the red shift at physiological temperature could be that it broadens the absorption window (70,71).

### 3.5 Target analysis

In addition to global analysis, we performed target analysis on 440 nm closed state TCSPC data at 279 K with the aim of resolving the rates of EET, charge separation (CS) and charge recombination (CR) for PSII. Figure 6 shows the kinetic models from target fitting, Figure 7 the species associated spectra (SAS) produced. Figure 8 shows how the concentration of different species evolves with time. Table 2 and Table 3 show the rates (in ns^-1^) and excitation vectors respectively. When the kinetic model is correct, the SAS resemble the spectrum of individual components, χ^2^ is close to 1 and residuals do not show any structure (Figure 7, Supplementary Fig. 11). From the SAS, the DAS can be reconstructed (Supplementary Fig. 14), showing agreement with our global analysis presented in Section 3.2. The excitation vectors of each component largely agree with the calculation obtained by global analysis of the number of pigments associated with each photosystem (∼1.76 : 1 ratio of PSII : PSI associated pigments, counting PSII antenna pigments as PSII-associated). Furthermore, if we estimate the relative absorption of FCP to PSII at 440 nm, based on FCP absorption as reported in (36) and the known pigment composition of PSII, a rough calculation gives approximately 22 FCP monomers per PSII dimer in total (Supplementary section IX). This corresponds to ∼352 pigments in the antenna ( ∼ 154 chl a, ∼ 154 fx, ∼ 44 chl c), and ∼94 pigments in the PSII dimer (∼74 chl a and ∼20 β-carotene).

**Figure 6.**
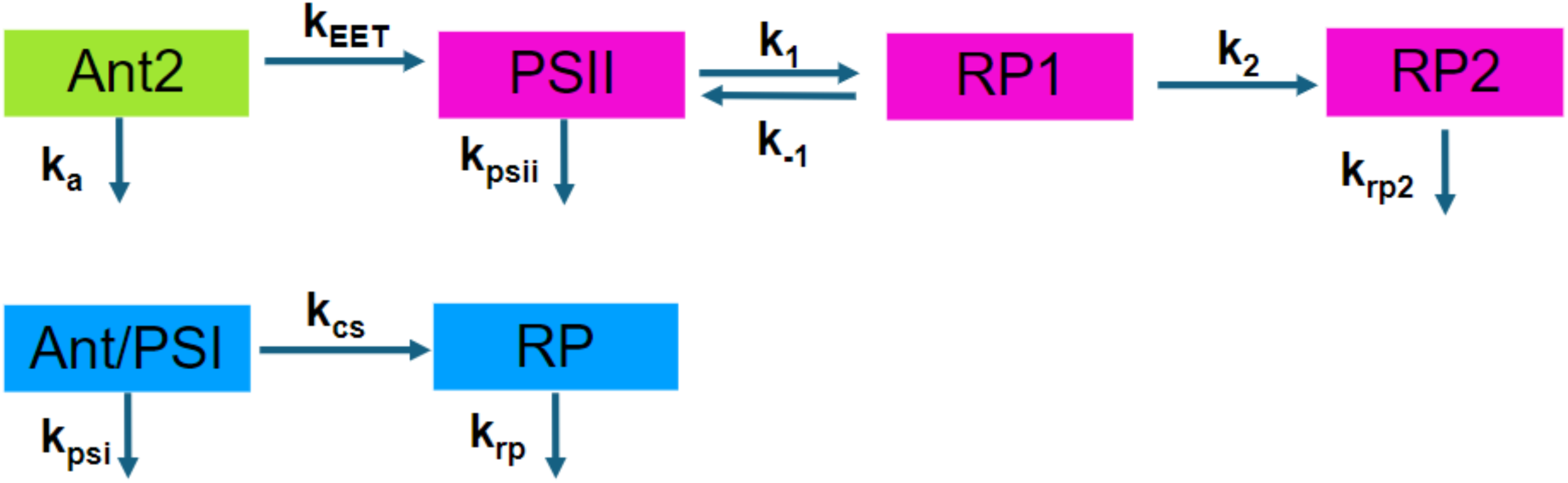
Target model for C. simplex obtained by analysis of TCSPC data at 279 K.

**Figure 7.**
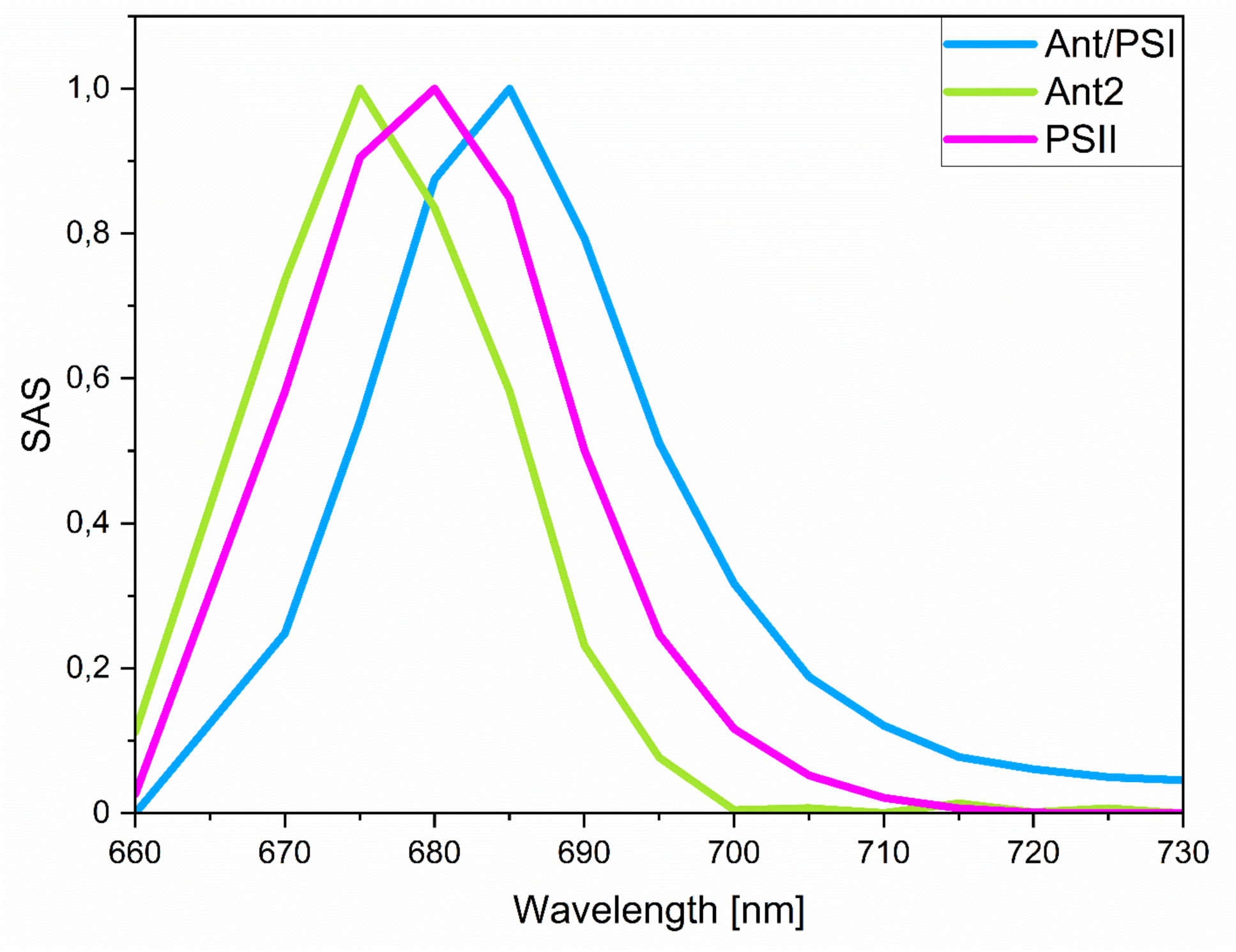
Species Associated Spectra (SAS) obtained from target analysis of closed state 440 nm TCSPC data at 279 K. Spectra are normalised to emission maximum for each species.

**Figure 8.**
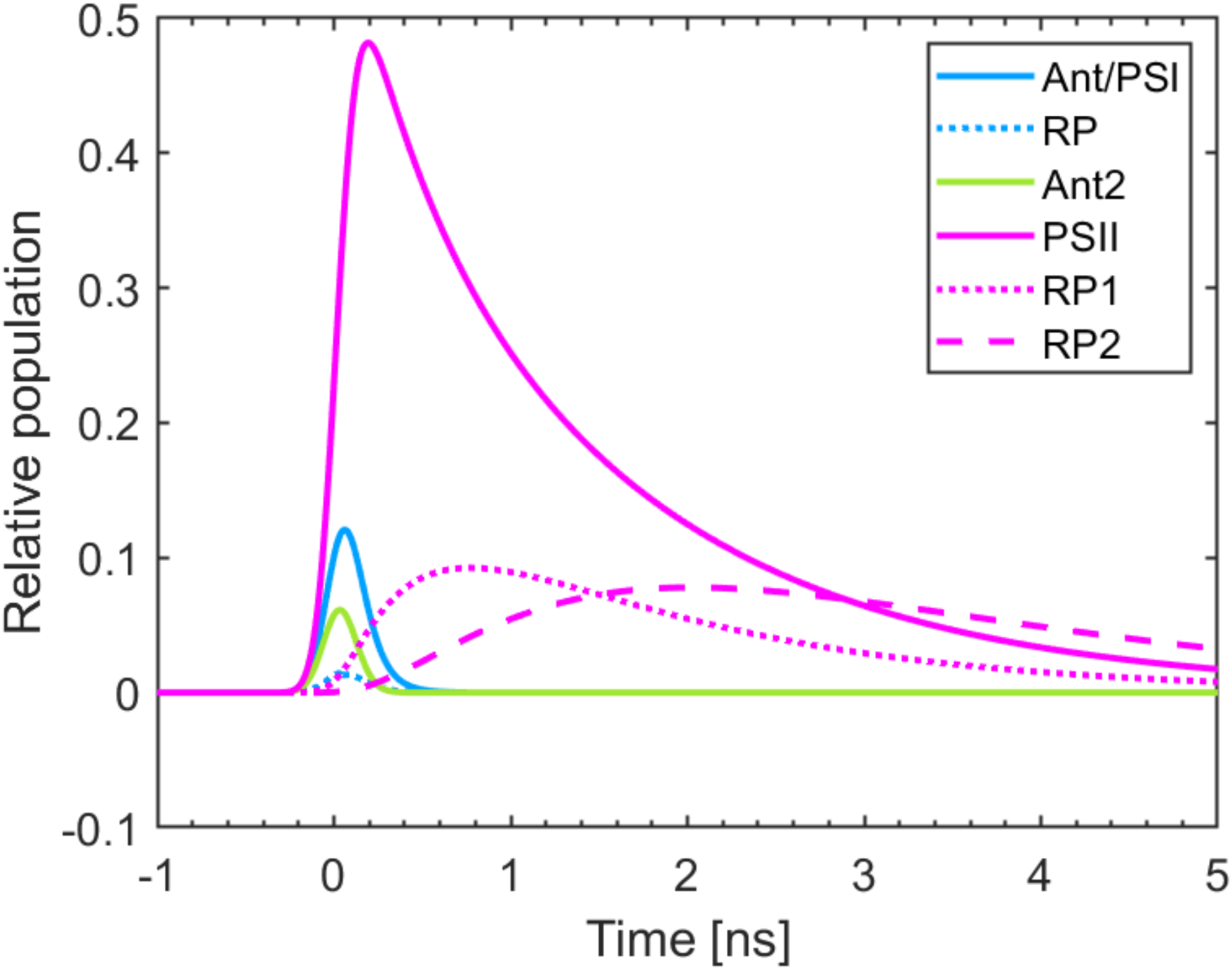
Concentration profile obtained from target analysis of closed state 440 nm TCSPC data at 279 K.

**Table 2.** Rates of EET and primary photochemistry for C. simplex obtained from target analysis. ^1^An upper limit of 30 ps was set based on the estimation of change in migration time from global analysis, in other words, migration was not allowed to be faster than 30 ps. ^2^A limit of 11 - 12 ns^-^1 was applied, corresponding to PSI decay of 90 ps – 83 ps, based on the results from global analysis (85 ps), since the PSI decay time is expected to be similar whether either global or target analysis is applied. ^3^The intrinsic decay lifetime of Ant2 was set to vary within the published decay times of isolated FCPs (20,109,110).

|  | Rate in ns<sup>-1</sup> |  | Rate fixed? |
| --- | --- | --- | --- |
| Ant2 | k<sub>a</sub> | 0.88 | Constrained<sup>3</sup> |
|  | k<sub>EET</sub> | 25.6 | Constrained<sup>1</sup> |
| PSII | k<sub>psii</sub> | 0.500 | Yes (104) |
|  | k<sub>1</sub> | 0.614 | No |
|  | k<sub>-1</sub> | 1.01 | No |
|  | k<sub>2</sub> | 0.988 | No |
|  | k<sub>rp2</sub> | 0.684 | No |
| PSI | k<sub>psi</sub> | 0.311 | Yes |
|  | k<sub>cs</sub> | 11 | Constrained<sup>2</sup> |
|  | k<sub>rp</sub> | 100 | Yes |

**Table 3.** Excitation vectors of target model at 440 nm excitation.

| Excitation vector (%) | 440 nm |
| --- | --- |
| Ant/PSI | 39 |
| Ant2 | 23.6 |
| PSII | 37.3 |

Similarly to other *in vivo* studies, we resolved one radical pair (RP) for PSI and two RPs for PSII (72–75). In addition, we resolved a separate component for PSII antenna (Ant2) transferring energy to PSII core. This allowed us to separate PSII antenna kinetics at 440 nm excitation, unlike other studies which bundle Ant2/PSII as a single compartment. One component representing PSI antenna and core combined (Ant/PSI) gives an adequate description of PSI kinetics. Since our global analysis indicated 4 degrees of freedom, we fixed several rates during fitting to avoid overparameterizing the model (Table 2). In particular, PSI radical pair kinetics are beyond the resolution of our setup. We therefore fixed the value of k_rp_ (PSI trapping) from the literature (100 ns^-1^ in several studies) (76).

From the kinetic model we obtained 3 SAS, which show a clear spectral separation between Ant2 (peaking at 675 nm), PSII (peaking at 680 nm), and Ant/PSI (peaking at 685 nm). The rate of EET from Ant2 to PSII was fast – 25.6 ns^-1^, corresponding to an EET lifetime of ∼40 ps. Recently, EET in PSII-FCPII supercomplex has been found to occur as quickly as 3 – 10 ps in the diatom *T. pseudonana* where 7 FCPs surround dimeric PSII and ∼20 ps in *C. gracilis* PSII-FCPII preparation where 22 FCPs surround each PSII dimer (46,77). *In vivo*, we observe the kinetics not only of tightly bound FCPIIs, but also of FCPs loosely associated with PSII. Therefore, our value of 40 ps is an averaged value across *all* PSII antenna. Our value (40 ps) is still much faster than that of LHCII to PSII in plant thylakoids when ∼8 LHC trimers are present per PSII dimer (∼150 ps), and is similar to the ∼35 ps migration time of BBY complexes, where 4.9 trimers of LHCII are present per PSII dimer (63).

In general, PSII in our models obeys the Exciton-Radical Pair Equilibrium (ERPE) hypothesis, where equilibration between excited states of antenna and RC occurs quickly (∼1.5 ps in the literature (78–80)), while the rate limiting step is the primary photochemistry, exhibiting reversible slow charge separation on a ns timescale, acting as a shallow trap (64,73–75,79). The ns-scale fluorescence seen would then reflect the different processes that RP1 can undergo; e.g. recombination back to the ground state, reformation of the excited state P680*, or population of the triplet state of the primary donor (81). The introduction of a back rate for the first apparent CS step in our model (k_- 1_) was necessary to achieve a good fit quality, and this back rate was found to be faster than the forward rate (k_1_). In other *in vivo* studies on closed-state diatoms, green algae, pine needles and isolated plant chloroplasts, k_-1_ is also faster than k1 (72–75). This situation also holds for isolated closed PSII membranes, where the k_-1_ (16 – 20 ns^-1^) was 4 times faster than the k_1_ (3.2 - 4.0 ns^-1^). This may be due to the large positive change in the Gibbs free energy ΔG° (∼ +40 meV) of charge separation caused by the presence of singly reduced Q_A-_, as proposed in reference (81). This in turn decreases the overall yield of charge separation in closed state, limiting the formation of ^3^P680 (triplet form of P680) by recombination, therefore limiting the production of reactive oxygen species (ROS) that can cause widespread cellular damage (81).

## 4. Discussion

Our most striking result is that *C. simplex* PSI is blue shifted at both 279 K and 77 K compared to other organisms. In Chukhutsina et al (40) at RT a component peaking at 690 nm was assigned to PSI. In contrast, *C. simplex* PSI peaks at 685 nm at 279 K. At 77 K, *C. simplex* PSI peaks at ∼700 nm, compared to *C. meneghiniana* where it peaks at ∼705 – 710 nm and has an additional ‘red chl’ peak at 717 nm (13). Isolated PSI cores in *C. gracilis* at 77 K emit around 700 – 705 nm while PSI-FCPI supercomplexes are even more red, peaking at 710 nm with a shoulder from 710 nm to 750 nm (82). In plants, isolated PSI core peaks around 730 – 740 nm at 77 K (83). Therefore, *C. simplex* PSI lacks the red chl sites seen in many other organisms, which fluoresce beyond 700 nm. Such red chls are important to PSI function in both the red and green lineage, encompassing higher plants, green algae, cyanobacteria and other diatoms. The red chls are involved in transferring energy uphill to bulk chls or P700^+^ (84,85) and have a photoprotective role in quenching excess excitation energy when P700 is reduced (86). Therefore, the lack of red chls in *C. simplex* is a significant result.

Unlike *C. simplex*, in *C. meneghiniana* and *P. tricornutum*, a single red compartment can transfer energy (reversibly) to the PSI RC, at rates determined to be around the same for both diatoms, 150 ns^-1^ forward transfer (74) (with back transfer of 150 – 180 ns^-1^). It is not known with certainty whether the red forms in these diatoms are located in the PSI core or in the antenna. In plants, the most well studied red forms are located in antenna subunits Lhca2/3 and Lhca1/4 where they form two heterodimers emitting at around 728.5 nm and 731.5 nm at 77 K (76,87,88). Pea plant PSI also has red states in the core: three chls emitting at 705, 715 and 722 nm, each having distinct decay kinetics, indicating that they are disperse and not coupled (88). The green algae *Chlamydomonas reinhardtii* (*C. reinhardtii*), however, lack these red forms in the core (89). Recently, a ‘primordial’ cyanobacterial PSI structure of *Gloeobacter violaceus* (*G. violaceus*) lacking red chls was published (44). Comparison of *G. violaceus* PSI structure with that of other cyanobacteria and eukaryotes showed that one chl dimer and one chl trimer are missing in *G. violaceus* PSI, named Low1 and Low2 and found on PsaA and PsaB respectively (44,90).

Low1 and Low2 are responsible for steady-state emission at around 723 nm and 730 nm in cyanobacteria. Low2 is missing in some cyanobacterial and eukaryotic PSIs, whereas Low1 is absent only in *G. violaceus*. Low2 is absent in all diatoms due to the absence of PsaG in diatoms, which is required for the formation of Low2 (44). Sequence comparison of Low1 gene regions shows that *G. violaceus* has a mutation Phe243, which replaces a His241 that coordinates binding of one of the red chls of the dimer (Chl1A). Phe243 prevents binding of Chl1A by steric hindrance. However, *C. simplex* sequence contains this conserved His residue as well as other conserved regions of Low1 (44) (Supplementary Fig. 15). Further comparison of *C. simplex* Low1 region with *C. neogracilis*, *C. meneghiniana*, *P. tricornutum* and *T. pseudonana* show that they all contain this conserved His residue (Supplementary Section X). We therefore tentatively propose that the red chl compartment needed to model *C. meneghiniana* and P. t*ricornutum* PSI kinetics in vivo in Miloslavina et al. (20) originates from a completely different site. In *C. gracilis,* the presence of FCPI red-shifts the emission of PSI-FCPI relative to PSI core, therefore, the red-most forms may be in the peripheral antenna of diatoms, not in the core, which may contain higher-energy red forms. These antenna-located red chls may be absent in *C. simplex*.

Despite the lack of red chl, PSI of *C. simplex* apparently has a long decay lifetime of ∼85 ps, longer than *in vivo* plant PSI-LHCI (50 – 81 ps) or PSI-LHCI-LHCII of *Zea mays* with three LHCII trimers (100ps) (20,91–94). In plants, the red chls in Lhca2/3 and Lhca1/4 have a much greater influence on the overall trapping kinetics of PSI than the effect of bulk antenna enlargement (76). While the slow energy transfer between red chls and other pigments lowers the ratio of charge separation lifetime to EET lifetime (τ_CS_/τ_mig_ ratio) of PSI, since τ_CS_ is still > τ_mig_, the kinetics remains trap limited (76). Average trapping time is increased in plant species whose PSI-LHCI emission is red-shifted compared to other plants, indicating they contain red forms of a lower energy level (e.g. 81 ps in *H. vulgare* compared to 61 ps in *N. tabacum*.) (91) The PSI lifetime of *C. simplex* is longer than that of *C. meneghiniana* cells at RT which is 75 ps (in low light of 20 µE) and 51 ps (in medium light of 40 µE) and shorter than that of *P. tricornutum* in medium light of 40 µE (100 ps) (20,40). Since *C. simplex* lacks the red chls, the long trapping time in this diatom must be due to a very large PSI-FCPI supercomplex. Analogously, *C. reinhardtii* PSI-LHCI has red forms that are higher in energy relative to plants, allowing it to expand antenna size without slowing down the average lifetime of this complex relative to plants (93). A large supercomplex in *C. simplex* may be an adaptation to maximize light harvesting in low light conditions (95). *C. meneghiniana* increases its antenna size when light intensity is decreased (40). This differs from plants, where antenna size of PSI is independent of growth light conditions (94). In addition, variation has been found in the number of FCPIs bound to PSI in diatoms, ranging from 5 to 24, with some FCPs able to detach in high light/quenched state (31,82,96,97). An absence of red forms could be a polar adaptation related to low temperature; for instance, in the Antarctic green alga *Chlamydomonas priscuii* (*C. priscuii*), the 77 K fluorescence emission spectrum of the single-celled form shows only minimal red PSI fluorescence at ∼ 711 – 713 nm, compared to the colonial palmelloid form induced by higher temperatures, which has increased red chl fluorescence (52).

The kinetics of PSII in *C. simplex* also appear slowed relative to *C. meneghiniana*. At physiological temperature, our open state average PSII lifetime of 480 ps is slightly longer than that for *C. meneghiniana* in low light growth conditions of 20 µE (448 ps) and medium light growth conditions of 40 µE (395 ps) and *P. tricornutum* in medium light growth conditions of 40 µE (432 ps) (40). This agrees with the longer Δτ_mig_ seen in *C. simplex* (30 – 50 ps vs 13 ps in *C. meneghiniana*). In closed state our average PSII lifetime of ∼1223 ps is also slightly longer than that of *C. meneghiniana* and *P. tricornutum* – 1170 ps and 1188 ps (20). This suggests that the PSII antenna of *C. simplex* is slightly bigger than that of temperate diatoms. Phytoplankton productivity in the Southern Ocean is co-limited by low light and low iron (98–100). Under such limiting conditions, Antarctic diatoms have higher oxygen use efficiency, and similar or higher photosynthetic iron use efficiency, compared to temperate diatoms (1). This is associated with a larger absorption cross section of both PSII and PSI, suggesting the presence of large antenna combined with a low number of cores (101). Large antenna maximize photon capture in light-limiting conditions, while the low number of cores (and probably also low PSI : PSII ratio (102)) reduces the demand for iron from PSI and the photosynthetic electron transport chain. In some studies, an inverse relationship has been observed between antenna size and efficiency of PSII in Southern Ocean phytoplankton (1,101,103). However, in our study *C. simplex* PSII quantum efficiency remains high despite the large antennae size (81%, calculated by the method used in (104)). Therefore, the large antenna size seen in *C. simplex* is likely an adaptive characteristic of polar diatoms in low light growth conditions (10 - 15 µE), while temperate species like *C. meneghiniana,* while having relatively smaller antenna show similar PSII quantum efficiencies (82% in Chukhutsina et al. and 84% in Miloslavina et al.) (20,40).

Using target modelling, we resolved the rate of excitation energy transfer from PSII antenna to PSII - 25.6 ns^-1^, corresponding to a migration time of ∼40 ps. In our model, only one PSII core compartment and one PSII antenna compartment were required to adequately fit PSII kinetics. Similarly, other *in vivo* experiments have found that only one PSII compartment is required (105). In contrast, different *in vivo* studies (72,73,75) have determined that two PSII pools are required to fully describe PSII kinetics, with each pool having different rates of charge separation and recombination. The physical meaning of this is that different cores experience different structural environments, leading to different excitonic behaviour. For example, some PSII are surrounded by bigger antenna than others, or are located in different regions of the chloroplast membrane (apressed / non-apressed) (106,107). Hence, our homogenous pool of PSII indicates that all PSII in *C. simplex* have approximately the same environment and effective antenna size. This agrees with the earlier results of Miloslavina et al (74) on *C. meneghiniana* and *P. tricornutum*.

In *C. simplex*, our 279 K data shows a clear PSI component of significant amplitude (33-42% of total area) over a wide range of antenna excitation. Therefore, a significant number of FCPs transfer energy to PSI, agreeing with our long PSI lifetime and absence of red forms. This contrasts sharply with C. *meneghiniana*, where at RT most of the antenna are connected to PSII and PSI fluorescence is hardly visible upon 535 nm excitation (40). Since global analysis is unable to resolve the rates of individual processes, the EET character of the fastest DAS from global analysis tend to reflect more than one process. In our case, we assigned the negative part of the fastest DAS to EET occurring between FCPs and from FCPs to (mainly) PSII. At 279 K, the lifetime of this DAS component is 17 – 30 ps. At 77 K, it is 16 – 19 ps, which resembles a similar process in *C. meneghiniana* taking place in 25 ps (13). At 77 K, Yokono et al (45) resolved a 25 ps component for EET from *C. meneghiniana* FCPs to *both* PSI and PSII. However, in target analysis we resolved the EET lifetime of a specific step – EET from FCPs to PSII core only, occurring in ∼40 ps. Therefore, the energy transfer occurring from one FCP to another FCP clearly occurs faster than 16 – 25 ps.

Lastly, we explored the roles of different fx in *C. simplex*, by selectively exciting fx-red and fx-blue using different excitation wavelengths. In doing so, we demonstrated that fx-red transfers more energy to PSII than fx-blue. The situation is similar in *C. meneghiniana*, where at 77 K both steady state and time resolved data indicated that fx-blue transfers more energy to PSI, whereas fx-red excitation mainly leads to transfer of energy to PSII (13). There is a possible role for fx-blue and fx-red in regulation of excitation energy transfer. Tselios et al (108) found that in *T. pseudonana*, at low light fx-blue and fx-red both transfer significant energy to chl a, and are also able to conduct energy transfer from fx-blue to fx-red. This transfer is reversibly inhibited in high light, where energy transfer from fx-red to chl a is negligible (41), due to reduced production of fx-red over the acclimation period of 4 days. We have demonstrated that the fx-red to PSII energy transfer occurs in *C. simplex* under low growth light conditions. If C. simplex behaves similarly to T. pseudonana, then less energy transfer from fx-red to PSII occurs in high light. This would suggest a possible role for fx-red and fx-blue in redirecting excitation energy away from PSII and towards PSI as a long-term acclimation strategy when light is in excess.

## 5. Conclusion

This study has elucidated the basic *in vivo* kinetics of the polar diatom *C. simplex*, which remains a fascinating organism for studying the enigma of polar photosynthesis. In light of the current difficulties in unifying structural and spectroscopic results from isolated complexes, this study can provide a view of energy transfer and charge separation in the intact cell.

We have demonstrated that the antenna of PSI and PSII in *C. simplex* are relatively larger than in the temperate diatom *C. meneghiniana*. In addition, PSI in *C. simplex* is blue-shifted in its fluorescence and unequivocally lacks red forms that fluoresce beyond 700 nm.

Our study has used low-light growth conditions and the dark-adapted state *in vivo* as a starting point for exploration. Further studies on the high light adapted state and/or quenched state could provide further insight into how this diatom adapts to the extreme environment of Antarctica.

