## Supplementary Material for "Antarctic photosynthesis: energy transfer and charge separation in the diatom *Chaetoceros Simplex*"

Contents

(i) Growth light spectrum
(ii) Absorbance titration for steady-state emission spectra, demonstrating that reabsorption did not affect the emission spectra.
(iii) Dual-PAM results showing that addition of DCMU and hydroxylamine closes PSII.
(iv) Experimental checks for TCSPC data to ensure that sample was stable during the experiment and 'open' state was maintained.
  a. Pinacyanol decay trace at the start and end of the experiment
  b. Power titration for 'open' state
  c. Sample decay trace at the start and end of the experiment
(v) Degree of variability in TCSPC results
(vi) Quality of fit data
  a. TCSPC data
  b. Streak data
  c. Target modelling
(vii) Total fluorescence spectrum and reconstructed steady-state spectrum of 77 K streak data
(viii) Decay Associated Spectra (DAS) reconstructed from target modelling of 279 K TCSPC data
(ix) Calculation of FCP : PSII ratio from target modelling excitation vectors
(x) Comparison of Low1 gene region of different organisms
(xi) 5-component fit of 77 K streak datasets at 550 nm excitation

I. Growth light spectrum

The light spectrum of the growth chamber was measured with a UV/Visible spectrophotometer (Jaz Spectrometer, Ocean Optics, USA). The resulting spectrum (Figure 1) shows the emission of the blue, green and red light-emitting diodes.

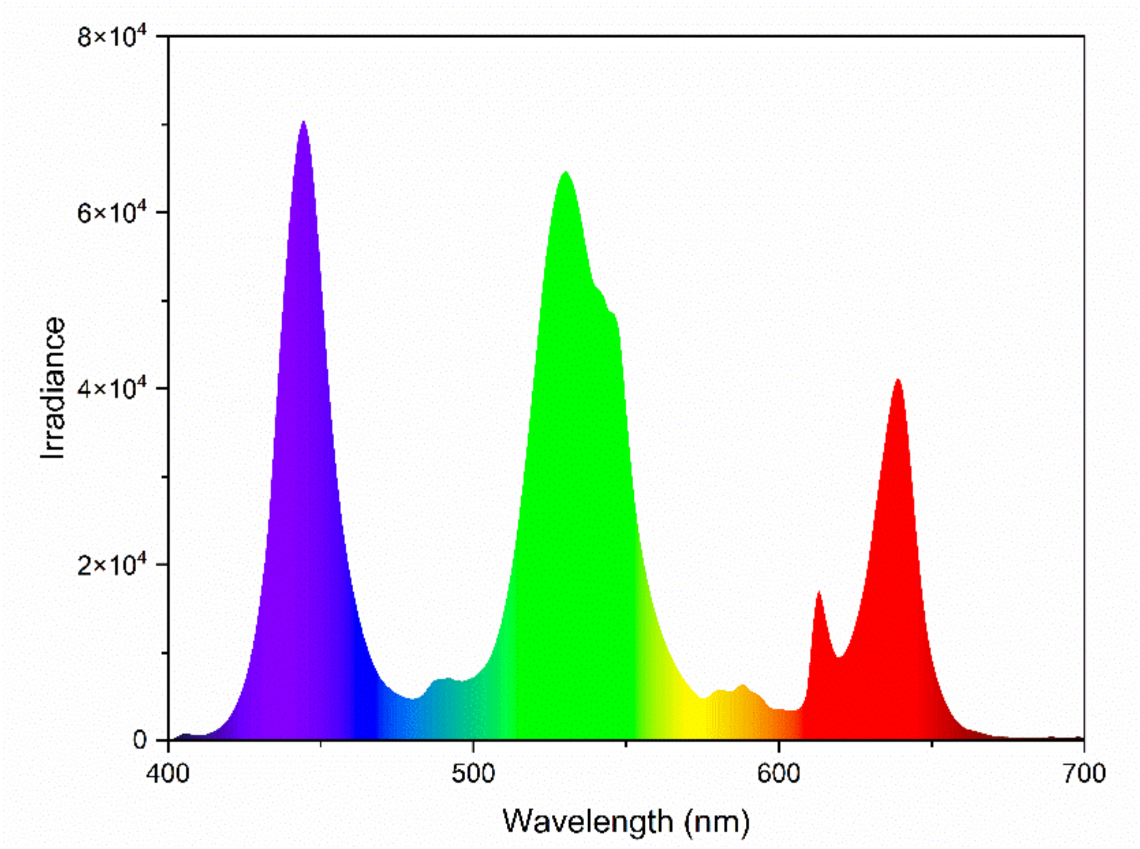


*Figure 1. Light spectrum of growth light (10-15 uE) with colour of visible light indicated.*

II. Absorbance titration for steady state emissions spectra

To characterize the effect of reabsorption on emission spectra, a titration series was done on samples of various absorbance values (0.1-0.5), of which an example is shown below for 279 K at 400 nm excitation (Figure 2). The presence of reabsorption is indicated by a red shift of the 680 nm peak and an enhancement of fluorescence above 700 nm. Our final experiments were performed on samples with absorbance of 0.1 or less.

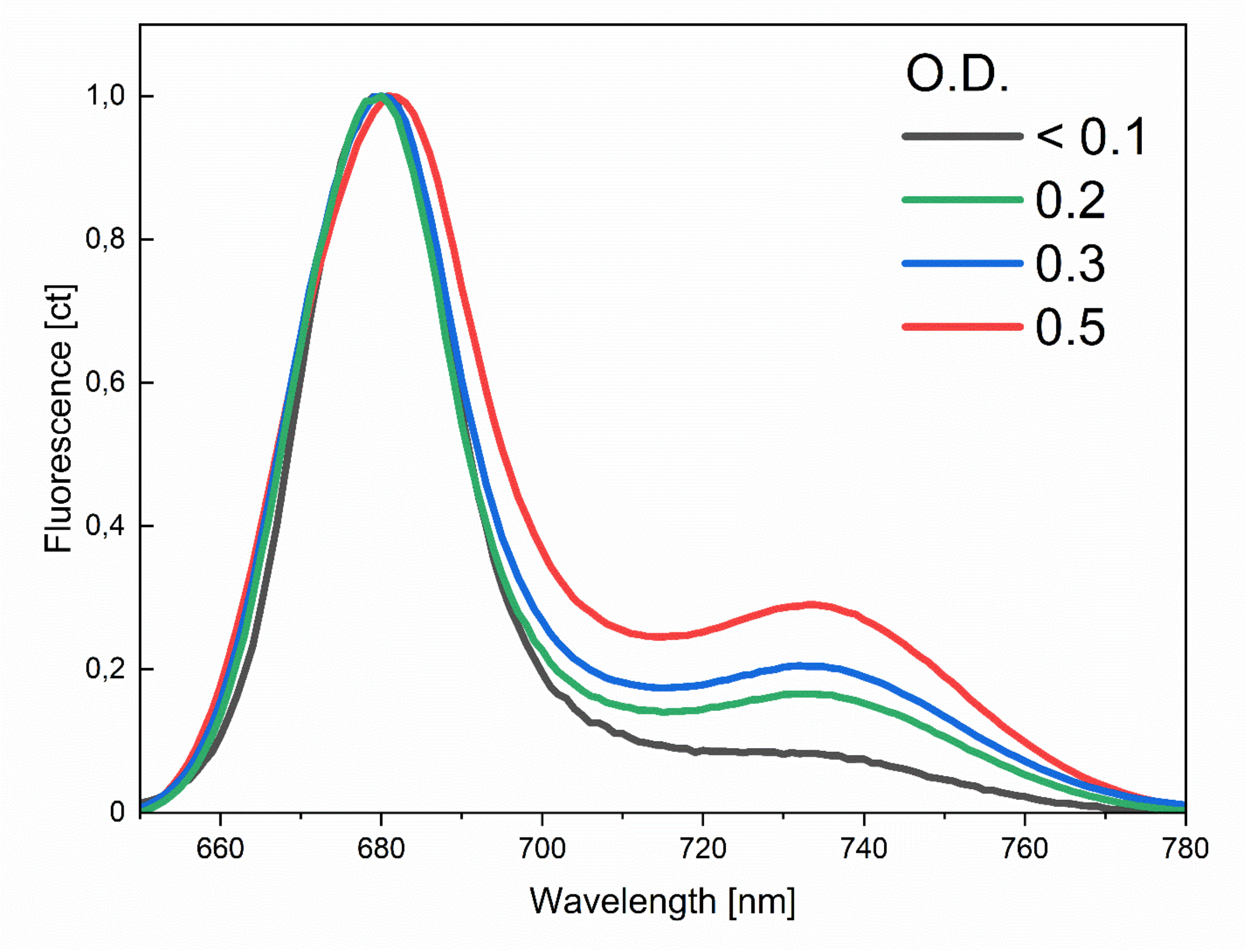


*Figure 2- Steady state emission of C. simplex samples of different absorbance (optical density, O.D.) and the effect of reabsorption.*

III. Dual-PAM results showing that addition of DCMU and hydroxylamine closes PSII.

To fully close PSII, the following protocol was followed (Figure 3): ‘open’ sample was loaded onto PAM (Dual-PAM 100, Walz, Germany) with a magnetic stirrer to constantly refresh the sample. Under constant measuring light of 24uE, a saturating pulse (SP) of 10,000uE was applied to measure the maximal fluorescence (Fm). After fluorescence returned to the $F_O$ (dark) level, cells were treated with 100uM DCMU and 1mM hydroxylamine (HA) at ~350s and the sample continued to be illuminated by 24uE light until $F_O$ rose to the level of $F_M$. Cells remained closed for 4-5 hours after this procedure.

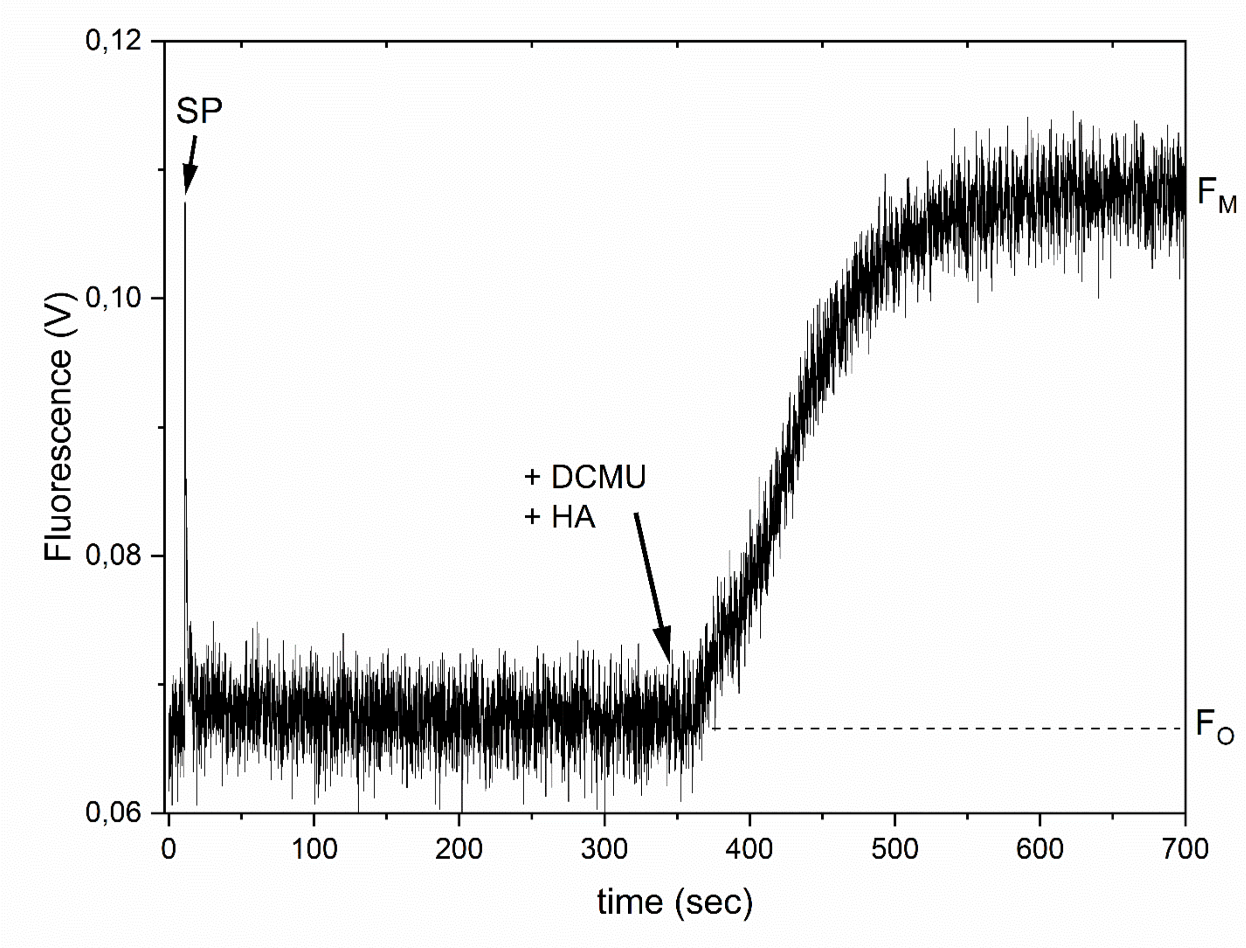


*Figure 3. Protocol for closing C. simplex cells with DCMU and hydroxylamine. SP: saturating pulse, DCMU: PSII inhibitor, HA: hydroxylamine, Fm: maximal fluorescence, Fo: dark fluorescence level*

IV. Experimental checks for TCSPC

Several data quality checks were applied to all TCSPC experiments. Firstly, the decay of pinacyanol in methanol was measured before and after the experiment to ensure that the instrument response function (IRF) drift was minimal (Figure 4). To ensure that the laser illumination or time elapsed did not change the sample, at the beginning and end of each experiment a decay trace was measured at 680 nm detection wavelength and compared to the original trace (Figure 5.). In addition, for 'open' state samples, power titrations were performed to ensure that sample was truly in the 'open' configuration (Figure 6).

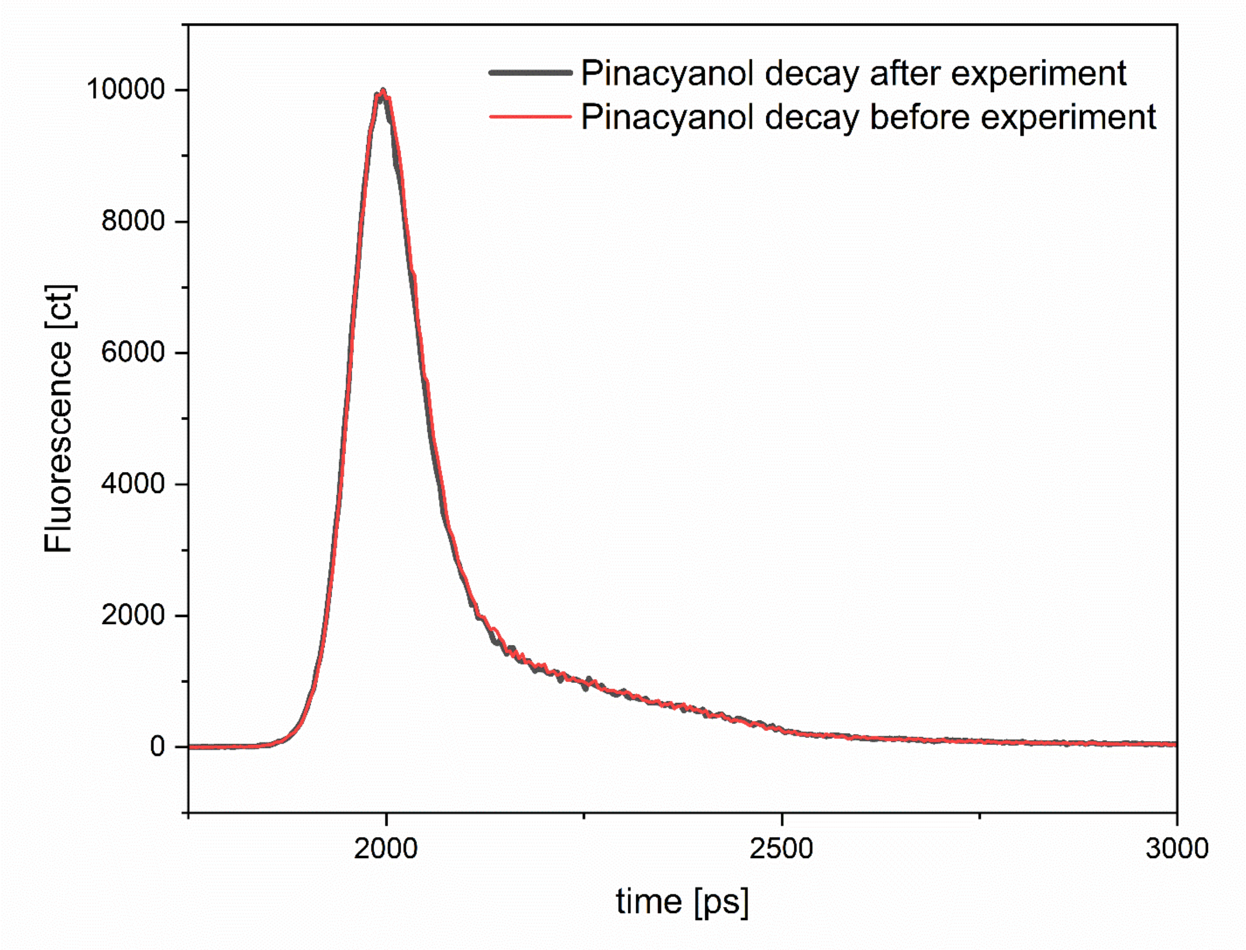


*Figure 4 – Decay of pinacyanol in methanol measured at 440 nm excitation and 680 nm detection before and after the experiment.*

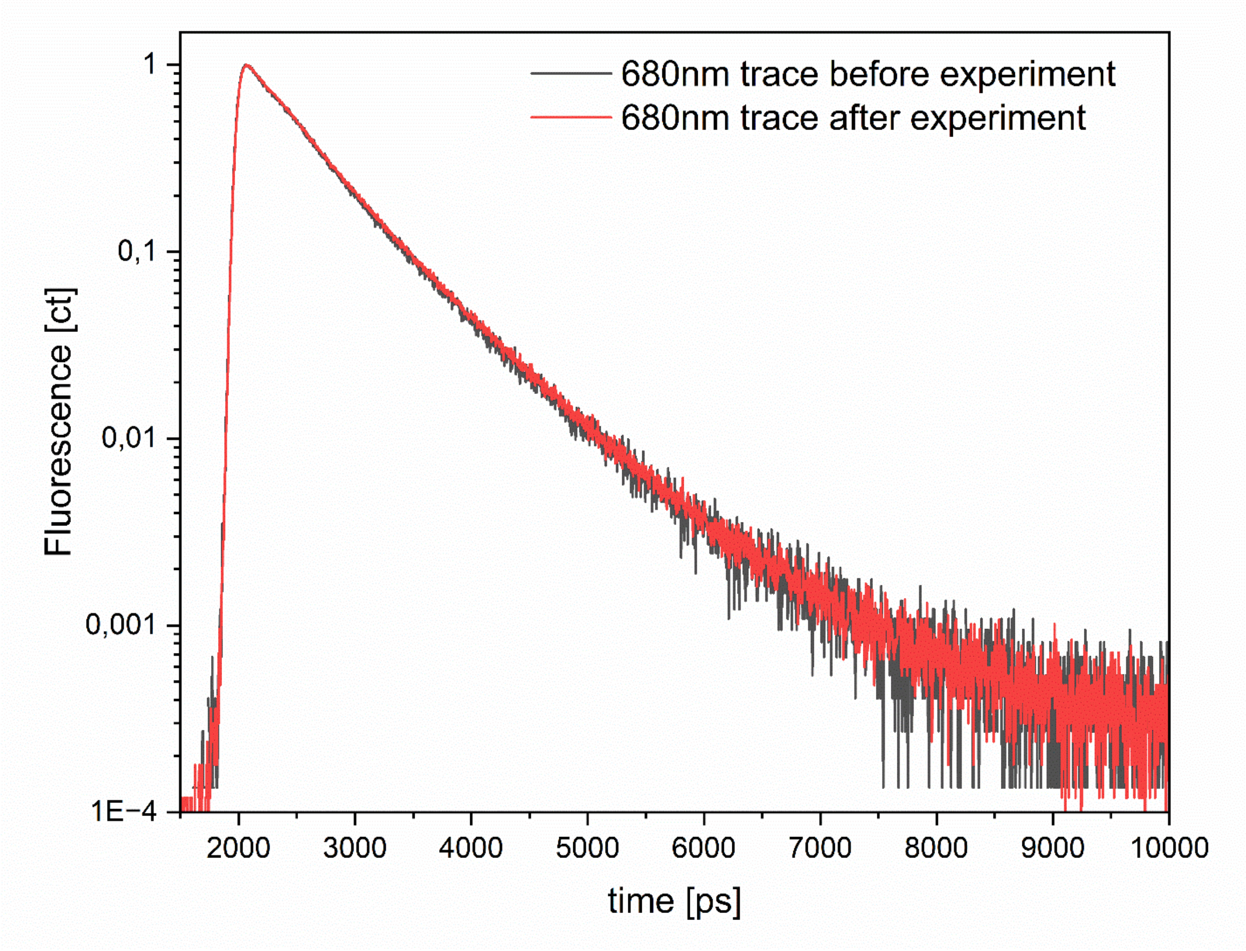


*Figure 5 – Example of 680 nm decay trace of an open state sample measured before and after data collection. The lifetime of the sample is the same in both cases.*

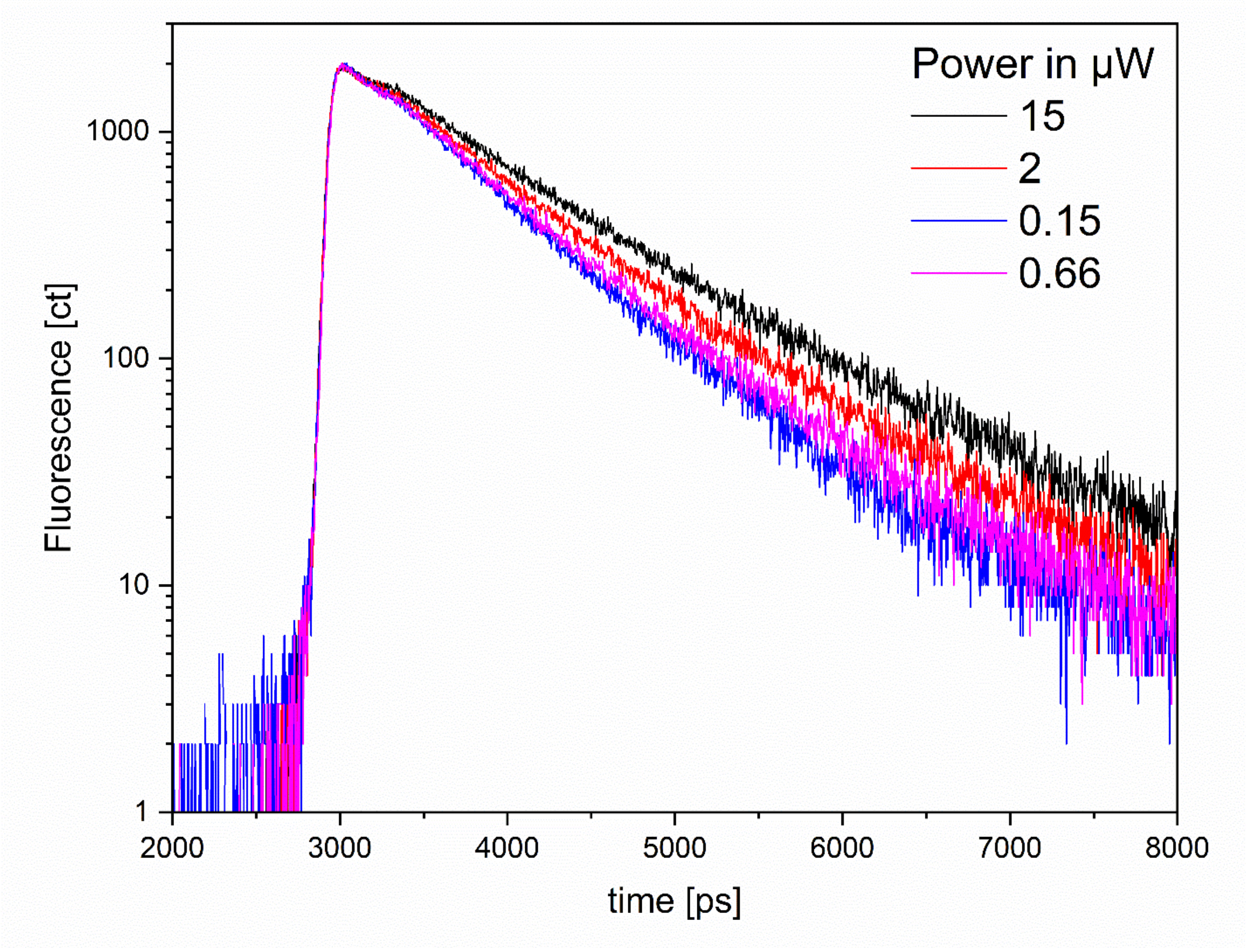


*Figure 6 – Power titration. Decay trace at 680 nm was measured for the open sample with different powers (shown here is an example at 440 nm excitation). Power was steadily decreased from 15uW to 0.66uW until no further shortening of the lifetime was seen, upon which the sample was deemed to have 'open' PSII.*

V. Degree of variability in TCSPC results

Decay curves from various experimental days were compared to one another, showing some sample variability, shown in (Figure 7) for cells in open state.

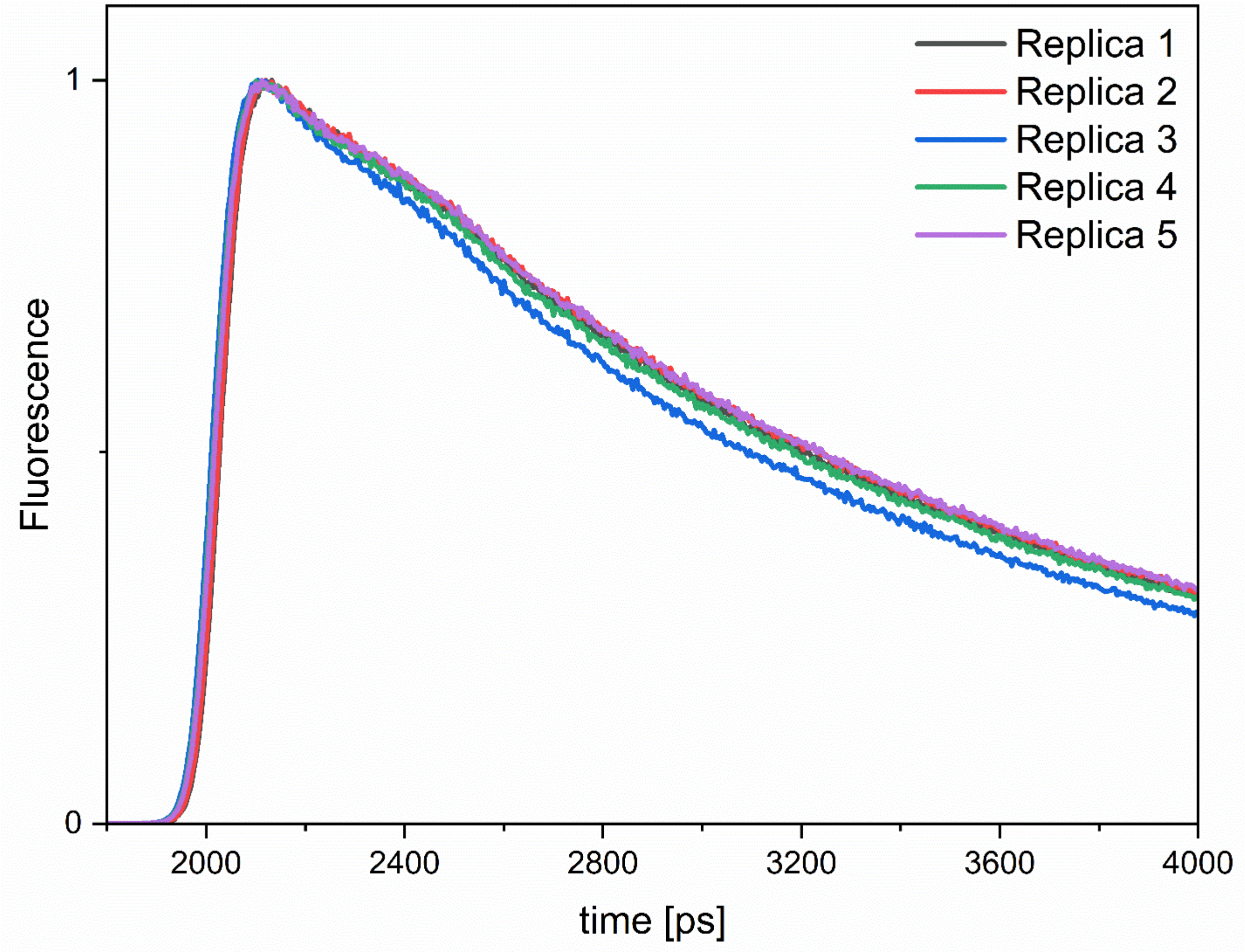


*Figure 7 – Decay trace of open samples measured on different days at 467 nm excitation and 680 nm detection, showing some sample variability as seen in the slightly different lifetime.*

This variability was reflected in the average lifetimes and average PSII lifetimes which differ by 10-30ps between replicates (cells harvested on different days).

VI. Quality of fit

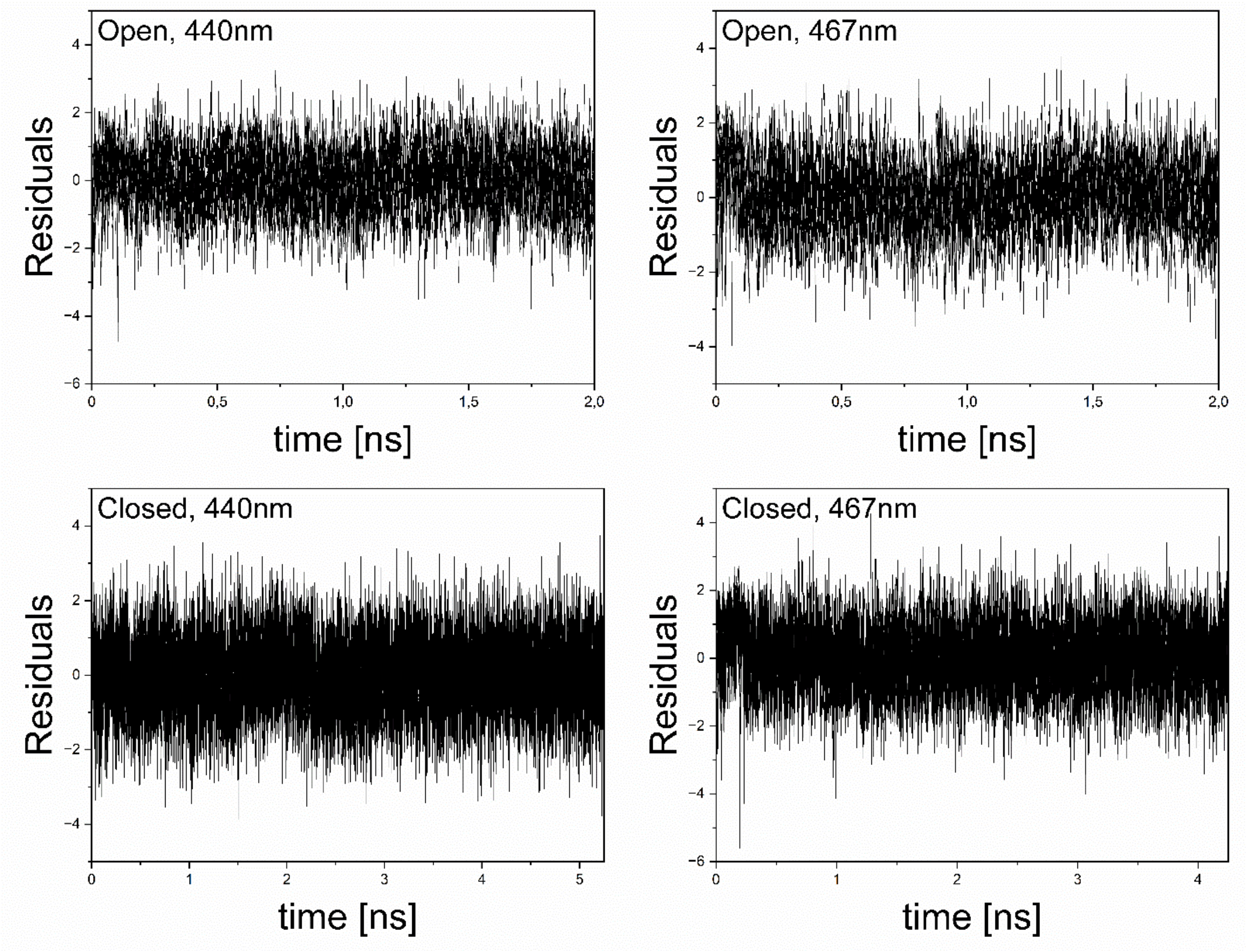


*Figure 8. Residuals from 279 K TCSPC data global fits. 1.0 < $\chi^2$ < 1.1 for all fits.*

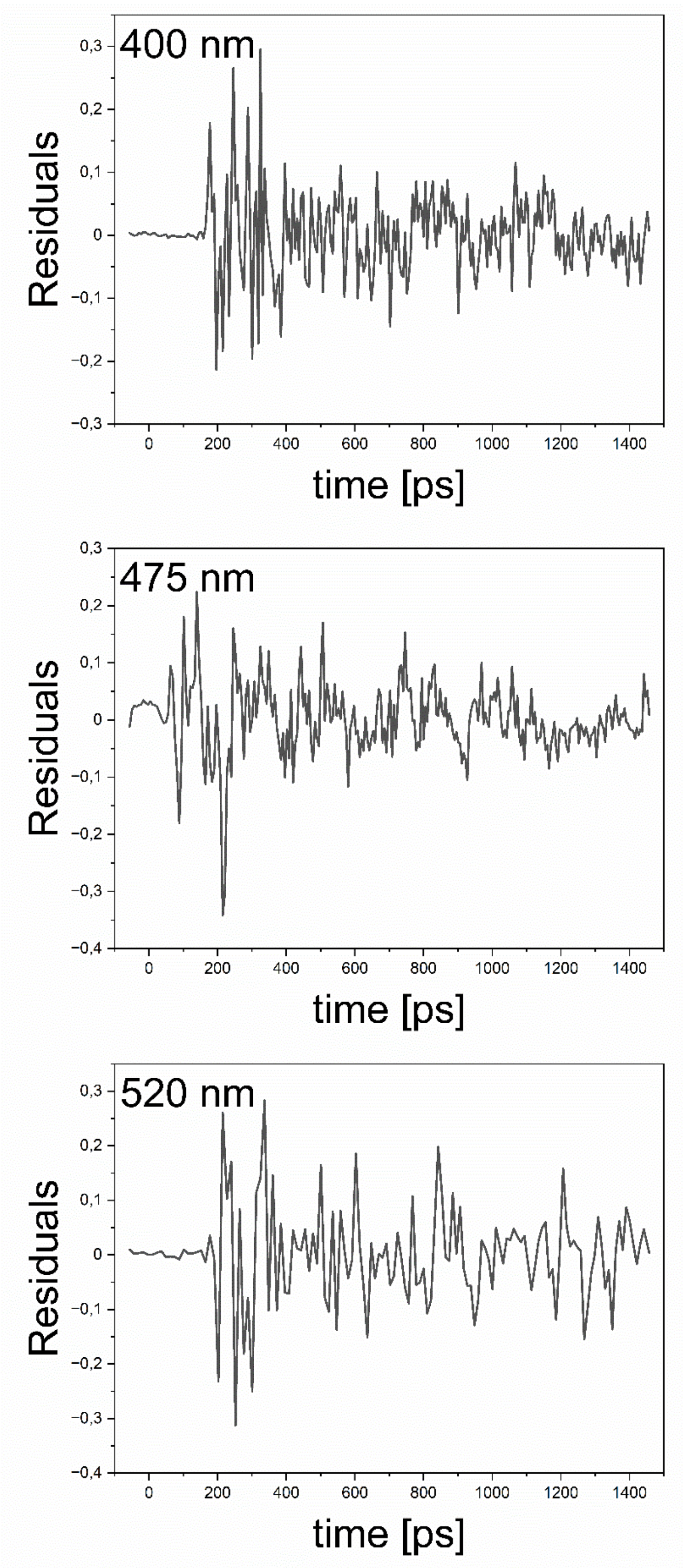


*Figure 9. First left singular vectors from 279 K streak data global fits for time range 4. Left singular vectors result from the singular value decomposition (SVD) of the residual matrix (1).*

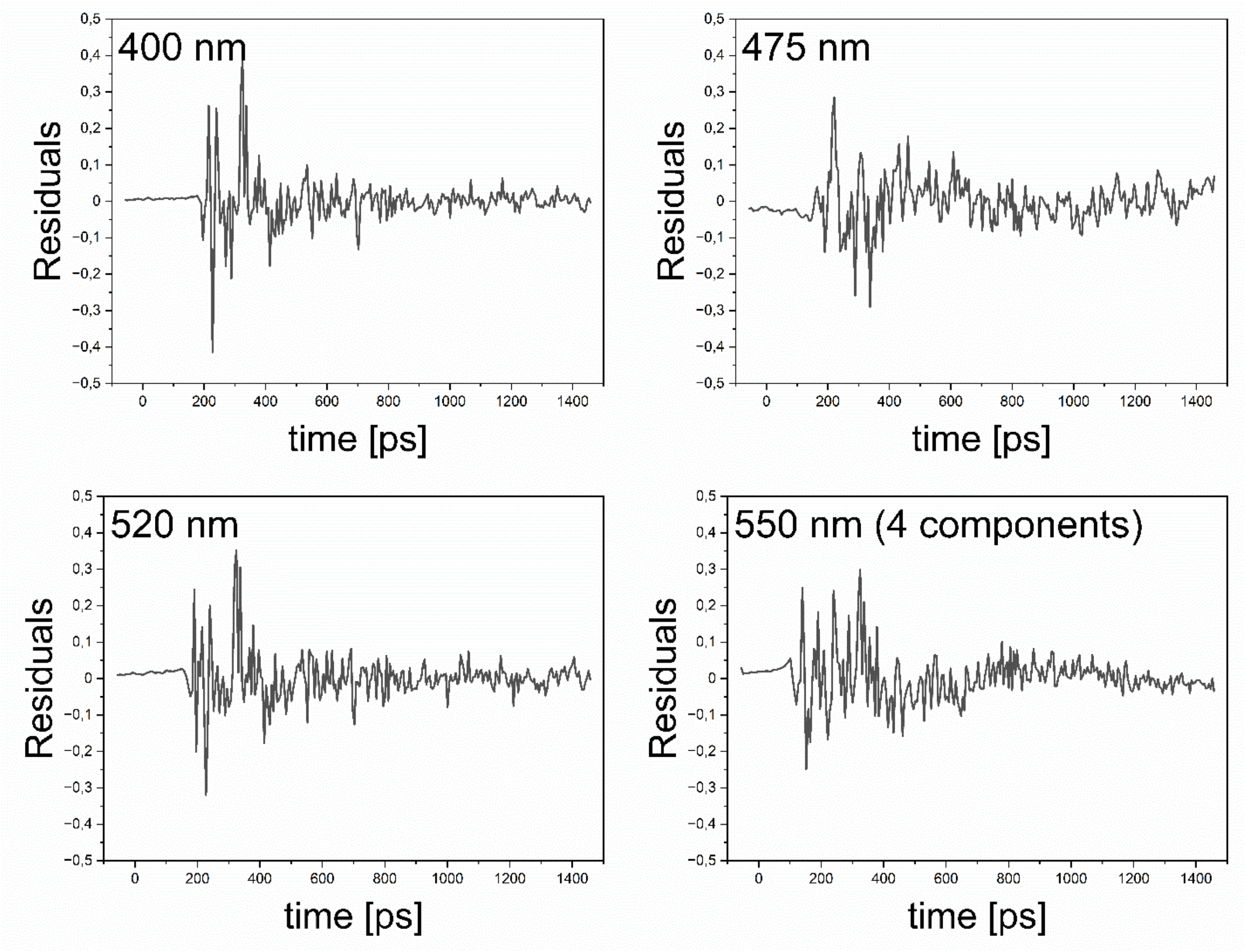


*Figure 10- First left singular vectors from 77 K streak data global fits for time range 4.*

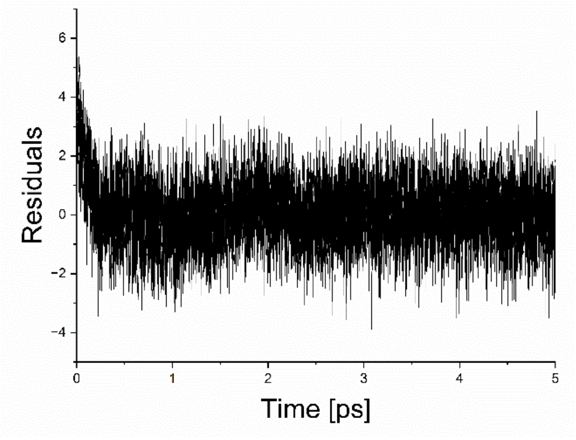


*Figure 11- Residuals from target modelling of TCSPC data.* $\chi^2 = 1.228$

VII. Reconstructed steady-state spectra and sum of DAS for 77 K streak data

The total steady-state fluorescence spectrum can be reconstructed from the global analysis results as the sum of each resolved DAS amplitude multiplied by the corresponding lifetime (Figure 12). The reconstructed spectrum (normalised to the maximum) is the same regardless of excitation wavelength, agreeing with our steady state results. The slight increase in red fluorescence at 475 nm excitation can be attributed to a slightly higher O.D. of that sample. Note that the wavelength resolution for streak (2.7 nm) is less than for our steady state setup (1 nm).

In addition, the total spectra at t=0 are equal to the sum of the various corresponding DAS for a particular excitation wavelength. Hence all DAS obtained from global analysis of datasets were summed and normalized in the maximum (Figure 13). The resulting spectra are the same regardless of the excitation wavelength.

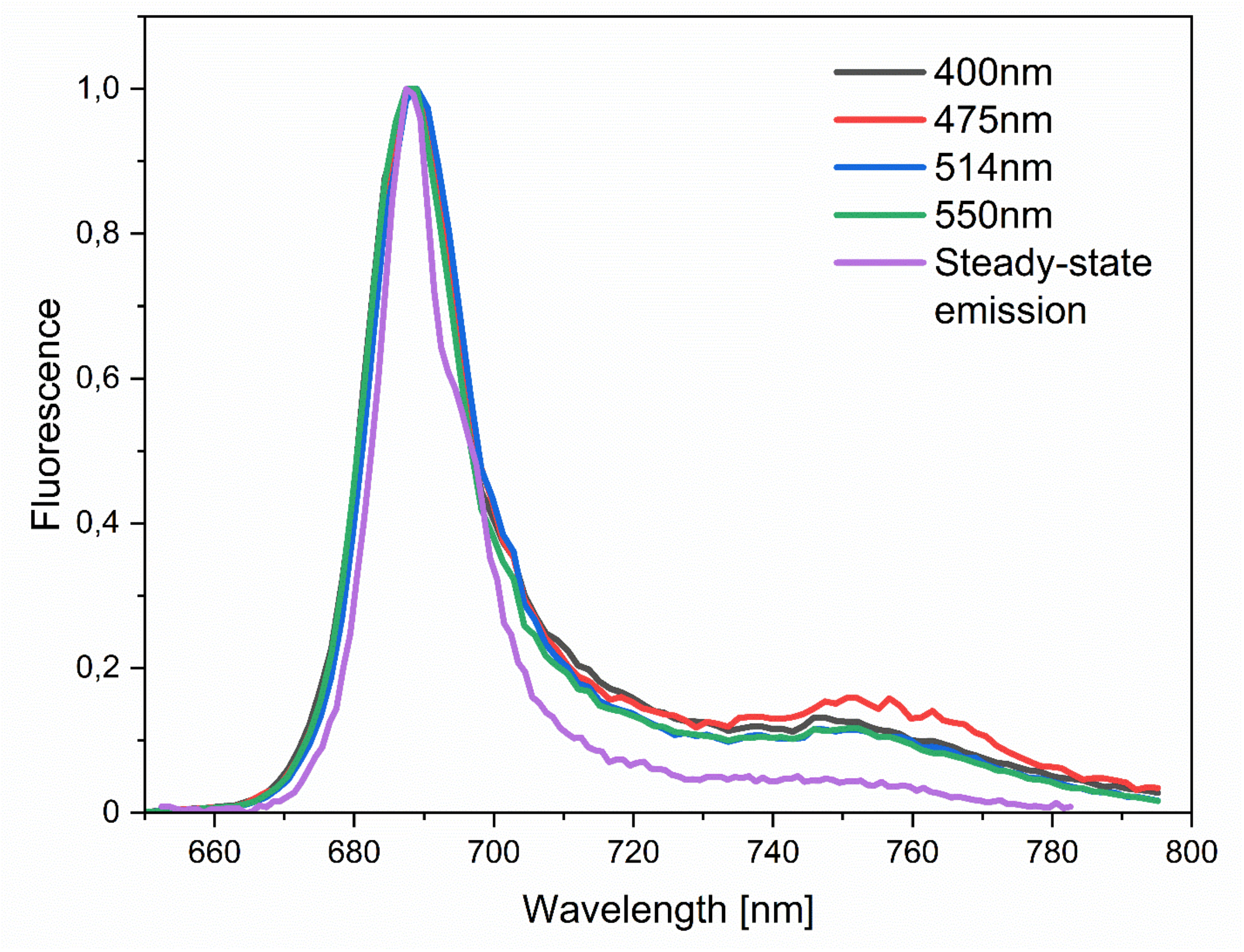


*Figure 12 – Reconstructed steady-state spectra of 77 K streak data, normalised to emission maximum. The steady-state emission spectrum at 77 K is included for comparison.*

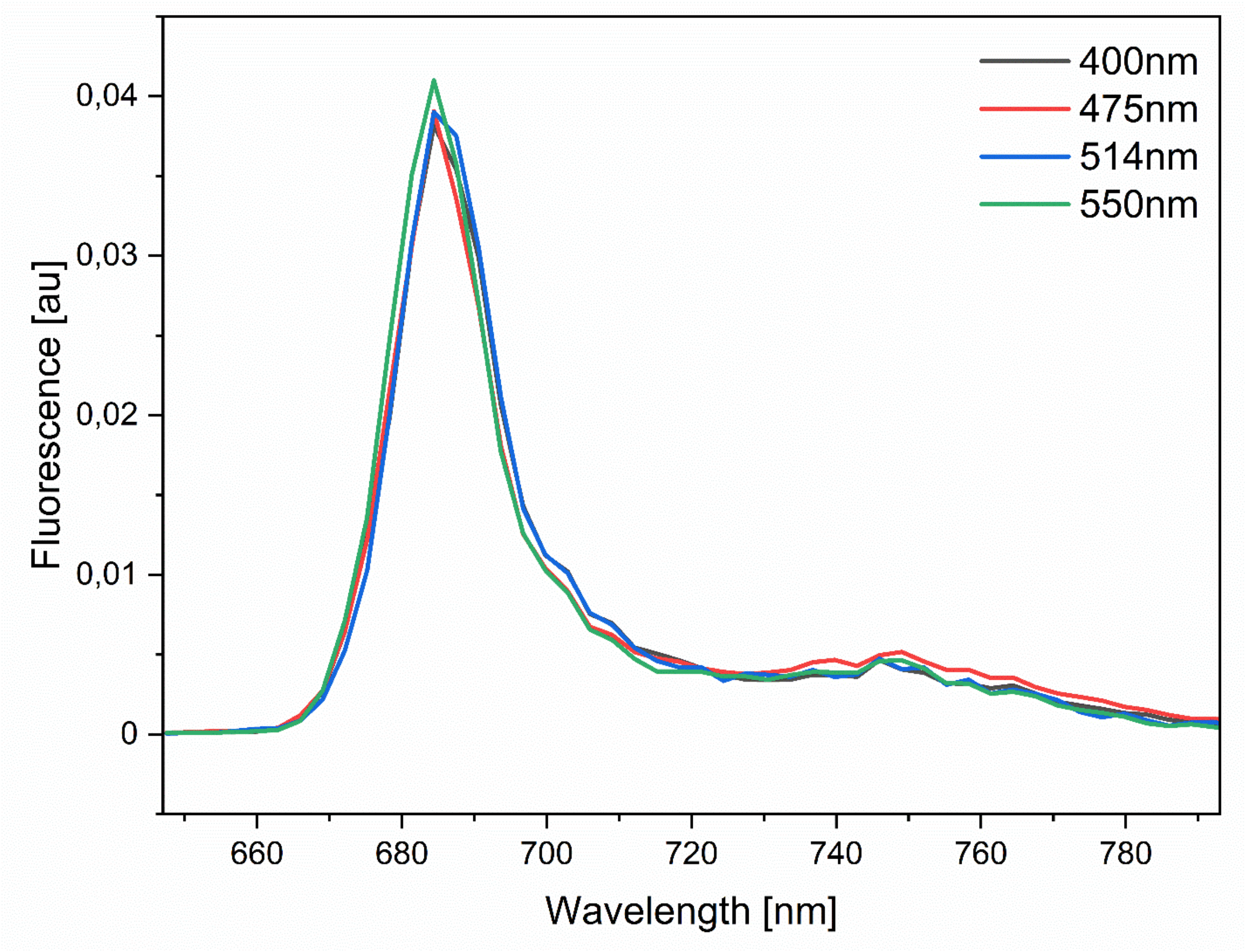


*Figure 13- Total fluorescence spectrum for 77 K streak data, normalised to emission maximum.*

### VIII. Decay Associated Spectra (DAS) reconstructed from target modelling of 279 K TCSPC data

Using the results of target modelling, the DAS of the closed state 440 nm sample was reconstructed (Figure 14), showing the energy transfer character of the ~40 ps component.

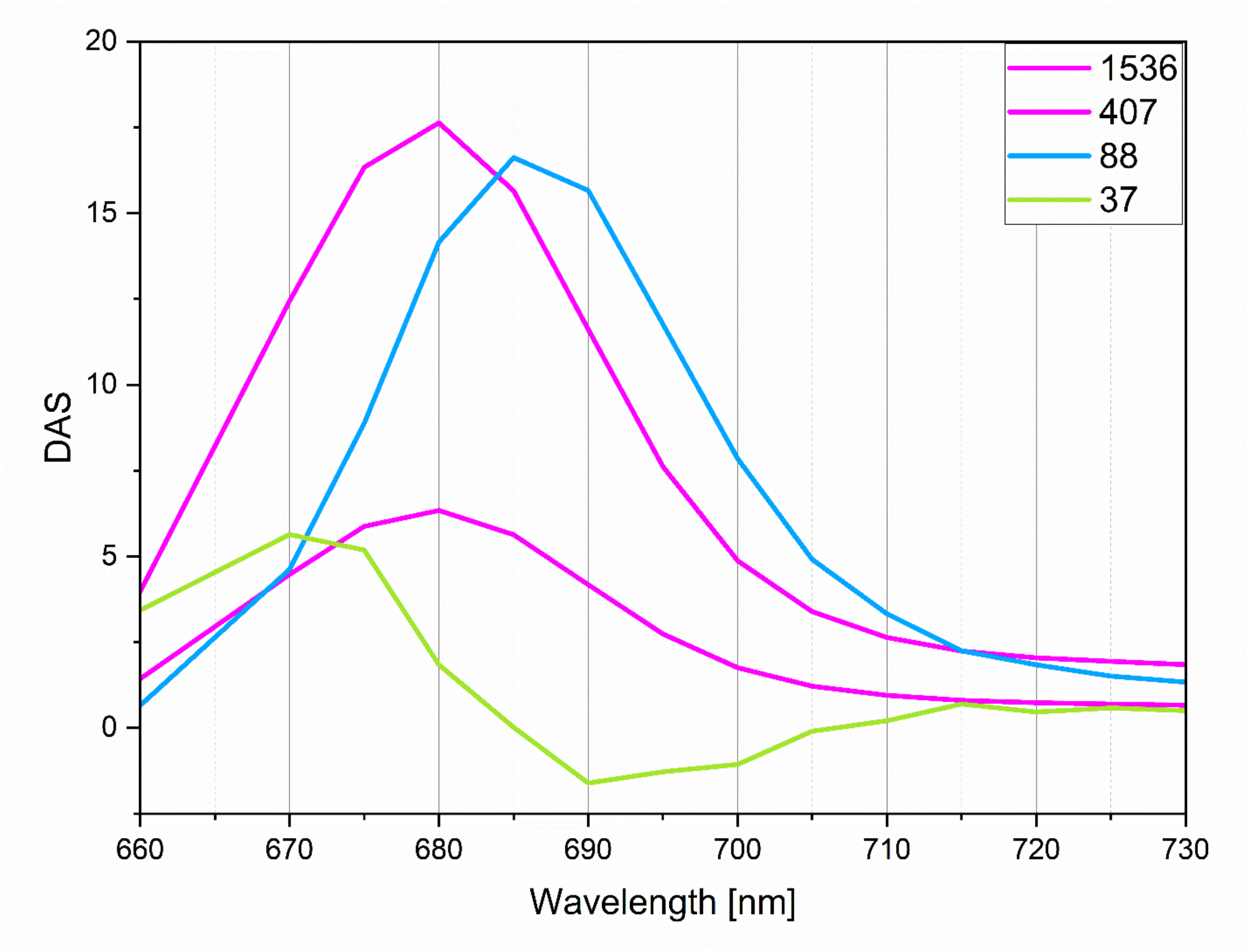


*Figure 14. Reconstructed DAS of 440 nm data. Lifetimes are given in ps. Legend: pink, PSII components, green, EET component, Blue, PSI component.*

### IX. Calculation of FCP : PSII ratio from target modelling excitation vectors

Since the excitation vectors obtained for each component are proportional to the effective absorption-cross section, using the relative absorption of individual pigments at that excitation wavelength, and the pigment composition of FCPs and PSII, we can find the ratio of FCP : PSII.

Premvardhan et al (2,3) reported the relative absorption of pigments (chl a, chl c, fx) for FCP trimer and oligomer from *C. meneghiniana*, measured in arbitrary units. Each FCP monomer contributes an absorbance of 0.175 (arbitrary units).

For PSII dimer, normalising to chl a, using the relative absorption of beta-carotene to chl a (4), and the ratio of 74 chl a: 20 beta carotene in PSII dimer, this gives 6.675 arbitrary units of absorbance.

Since the ratio of excitation vectors of PSII antenna : PSII dimer is 1 : 1.76,

$$\frac{FCP}{PSII\ (dimer)} = \frac{\left(\frac{1}{0.175}\right)}{\left(\frac{1.76}{6.675}\right)} = \sim 22$$

### X. Comparison of Low1 gene regions between different organisms

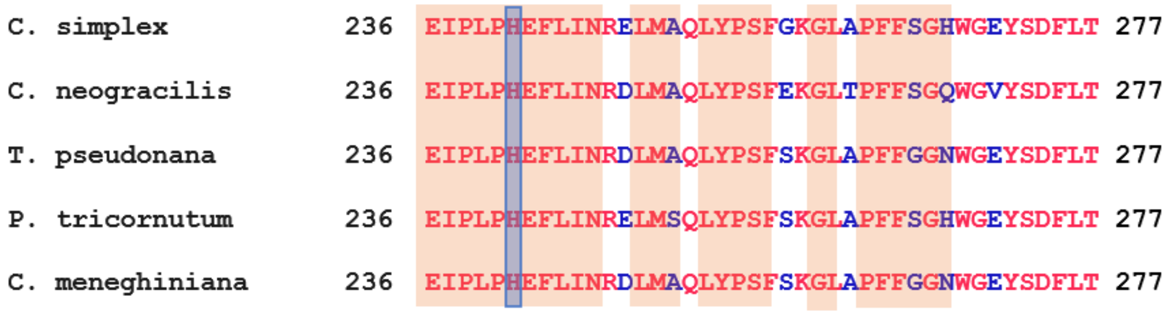


*Figure 15. Sequence comparison of Low1 gene regions of the diatoms C. simplex, C. neogracilis, T. pseudonana, P. tricornutum and C. meneghiniana. Amino acid identities are in red font and any amino acid differences detected during alignment are in blue font. Conserved regions as noted in Kato et al. (5) are highlighted in orange boxes, and the His241 residue is highlighted in a blue box.*

XI. 5-component fit of 550 nm 77 K streak dataset

For the 550 nm dataset, an additional component was added, producing a 5-component DAS fit (Figure 16), and this improved the residuals substantially (Figure 17). An interpretation of the physical meaning of the components is presented here.

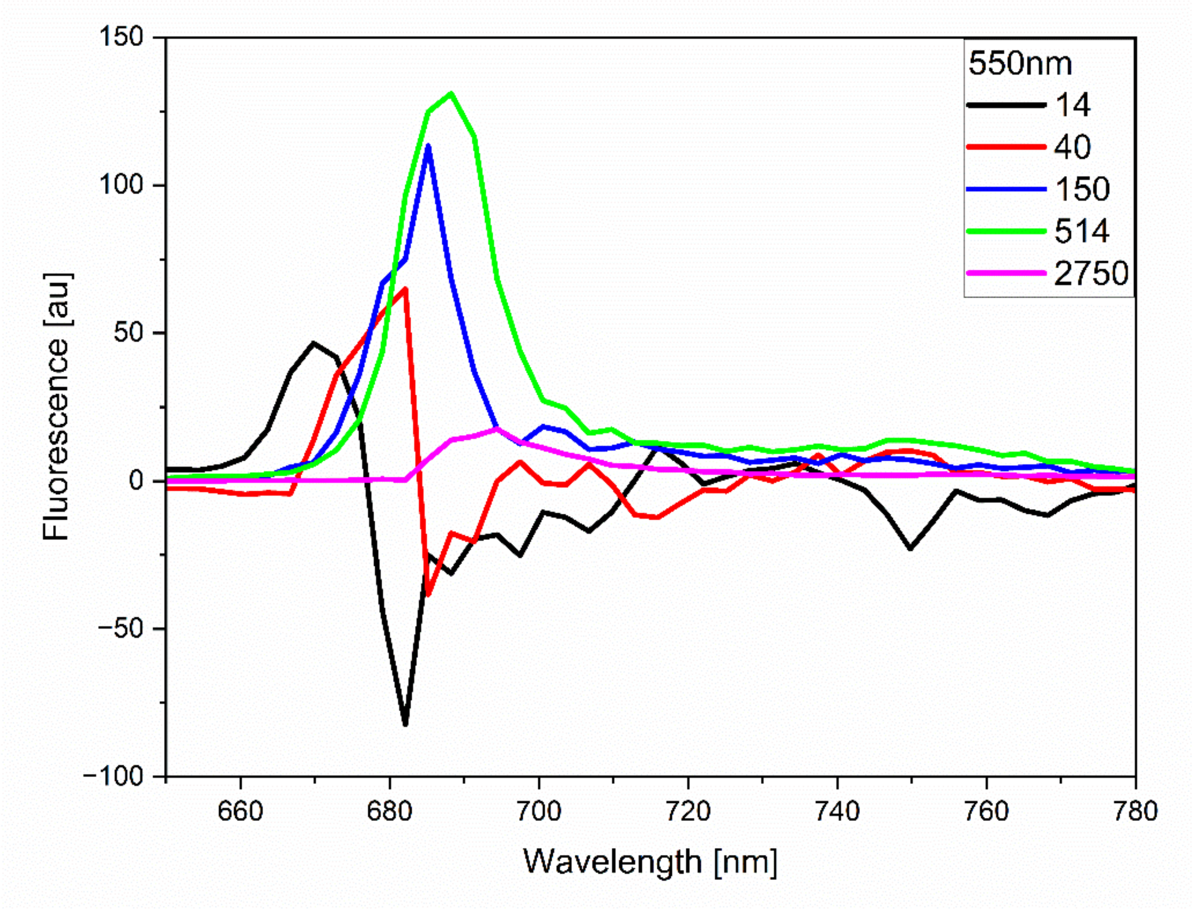


*Figure 16- 5-component DAS fit of 77 K streak data at 550 nm excitation. Lifetimes are given in ps.*

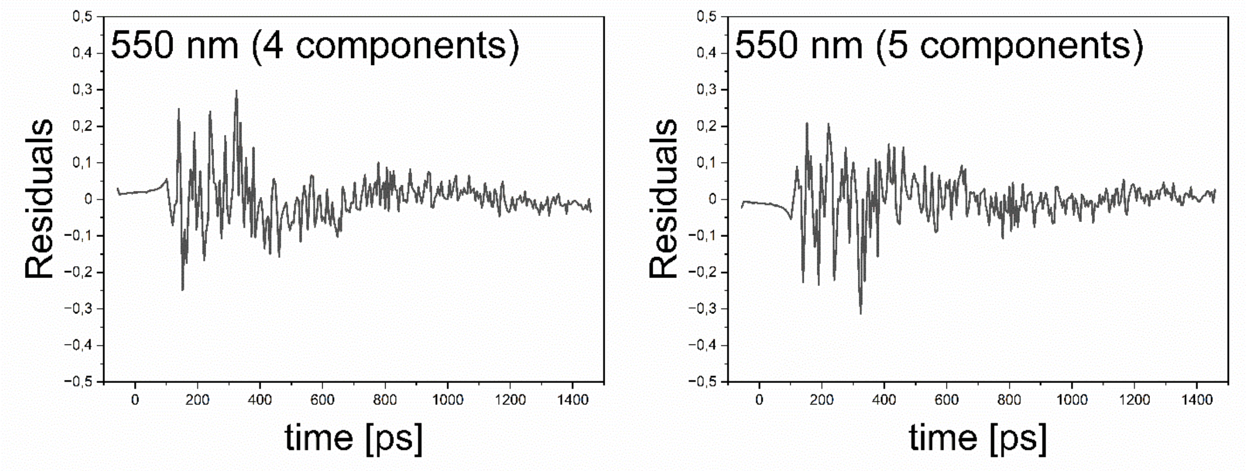


*Figure 17- Residuals from 5-component fit of 77 K streak dataset at 550 nm excitation*

In the 5 component fit, two populations of FCPs are revealed- one emitting at 670 nm (1st component, black) and transferring within ~14 ps to a FCP species emitting at a maximum of 682 nm (2nd component, red) that transfers energy to other species at a timescale of ~40 ps (negative peak at 685 nm). The third component (blue, 150 ps) emits with a main peak at 685 nm and could be due to a fast PSII decay (emission maximum 685 nm) combined with PSI decay (shoulder at 700 nm). As expected, the amplitude of PSI decay at 700 nm is very low, since very little energy transfer from FCP to PSI occurs at 550 nm. The fourth component (green, 514 ps) is mostly due to PSII decay. Lastly, the fifth component (magenta, ~2.7 ns) resembles the spectrum of aggregated FCP;  note that this long ns lifetime is not well resolved by the time window of the streak camera.
